# Meditation attenuates Default-mode activity: a pilot study using ultra-high strength MRI

**DOI:** 10.1101/2023.01.02.522524

**Authors:** Saampras Ganesan, Bradford Moffat, Nicholas T. Van Dam, Valentina Lorenzetti, Andrew Zalesky

**Affiliations:** Melbourne Neuropsychiatry Centre, Carlton, Victoria 3053, Australia; Department of Biomedical Engineering, The University of Melbourne, Carlton, Victoria 3053, Australia; Melbourne Brain Centre Imaging Unit, Department of Radiology, The University of Melbourne, Parkville, Victoria 3052, Australia; Neuroscience of Addiction and Mental Health Program, Healthy Brain and Mind Research Centre, School of Behavioral and Health Sciences, Faculty of Health, Australian Catholic University, Fitzroy, Victoria 3065, Australia; Melbourne School of Psychological Sciences, The University of Melbourne, Melbourne, Victoria 3010, Australia

## Abstract

**Objectives:** Mapping the neurobiology of meditation using 3 Tesla functional MRI (fMRI) has burgeoned recently. However, limitations in signal quality and neuroanatomical resolution have impacted reliability and precision of extant findings. Although ultra-high strength 7 Tesla MRI overcomes these limitations, investigation of meditation using 7 Tesla fMRI is still in its infancy.

**Methods:** In this feasibility study, we scanned 10 individuals who were beginner meditators using 7 Tesla fMRI while they performed focused attention meditation and non-focused rest. We also measured and adjusted the fMRI signal for key physiological differences between meditation and rest. Finally, we explored the 2-week impact of the single fMRI meditation session on mindfulness, anxiety and focused attention attributes.

**Results:** Group-level task fMRI analyses revealed significant reductions in activity during meditation relative to rest in Default-mode network hubs, i.e., antero-medial prefrontal and posterior cingulate cortices, precuneus, as well as visual and thalamic regions. These findings survived stringent statistical corrections for fluctuations in physiological responses which demonstrated significant differences (p < 0.05/n, Bonferroni controlled) between meditation and rest. Compared to baseline, State Mindfulness Scale (SMS) scores were significantly elevated (F = 8.16, p<0.05/n, Bonferroni controlled) following the fMRI meditation session, and were closely maintained at 2-week follow up.

**Conclusions:** This pilot study establishes the feasibility and utility of investigating focused attention meditation using ultra-high strength (7 Tesla) fMRI, by supporting widespread evidence that focused attention meditation attenuates Default-mode activity responsible for self-referential processing. Future functional neuroimaging studies of meditation should control for physiological confounds and include behavioural assessments.

## 1 Introduction

The past decade has seen a noticeable expansion in research investigating the functional brain mechanisms underlying meditation (Ganesan et al., 2022a; Melis et al., 2022; Sezer et al., 2022; Young et al., 2018). Among various techniques, the fundamental practice of focused attention meditation is widely investigated in the scientific literature (Bishop et al., 2004; Ganesan et al., 2022a; Lutz et al., 2008). This technique entails focusing and sustaining attention on an object or experience (e.g., breathing sensations) in the present moment while actively noticing and disengaging from distractions (e.g., mind-wandering). Focused attention on bodily experiences (e.g., breath sensations) is also categorized as body-centred meditation within an embodied framework that incorporates the influence of bodily states on psychological processes (see Matko and Sedlmeier (2019) for detailed categorization). Focused attention meditation trains attentional capacity, meta-awareness and interoceptive sensitivity necessary to effectively approach other advanced meditation states and techniques (Jha et al., 2007; Laukkonen & Slagter, 2021; Trungpa, 2002; Valentine & Sweet, 1999). On its own, practice of this meditation technique can enhance attentional regulation and mitigate habitual thinking patterns that may be maladaptive (Laukkonen & Slagter, 2021; Wenk-Sormaz, 2005). Owing to the benefits (Creswell, 2017; Galante et al., 2021; Shapiro & Walsh, 2003) and adverse events (Farias et al., 2020; Schlosser et al., 2019; Van Dam et al., 2018) associated with different meditation practices, understanding the neurobiology of specific meditation techniques can potentially complement self-report measures of meditation expertise, progress and outcomes. Additionally, reliable neural markers of meditation could potentially facilitate non-invasive neuromodulation therapeutics to assist psychiatric patients in practicing meditation.

With the advent of functional Magnetic Resonance Imaging (fMRI) technology, many studies have endeavoured to map the precise neural markers of meditation. Focused attention meditation (with breath or body sensations) is particularly amenable to fMRI investigations due to its simplicity, accessibility to beginner meditators, and significance across various traditions (Ganesan et al., 2022a; Matko et al., 2021). Conventionally, studies have used MRI scanners with magnetic strengths of up to 3 Tesla to examine the fMRI blood oxygen-level dependent (BOLD) processes underlying meditative states. This body of work has provided unique insights into the brain areas and brain networks that are frequently implicated by meditation. For example, most qualitative (Brandmeyer & Delorme, 2021; Feruglio et al., 2021; Laukkonen & Slagter, 2021) and quantitative (Fox et al., 2016; Ganesan et al., 2022a) neuroimaging reviews thus far have consistently highlighted that focused attention meditation is associated with reduced activity in the brain network ascribed to self-referential processing and mind-wandering, i.e., Default-mode network. Mitigation of distraction caused by spontaneous thought and mind-wandering is a core mechanism that facilitates sustained attention on the object of focus (e.g., breathing sensations) during focused attention meditation. However, compared to large-scale brain networks like Default-mode, Salience and Executive Control networks, evidence regarding involvement of neuroanatomically specific brain regions within these brain networks is weaker in focused attention meditation. For instance, 85% of the focused attention meditation literature consistently finds involvement by various regions that together constitute the Default-mode network. However, when considered separately, the network’s neuroanatomically specific constituent regions (e.g., posterior cingulate cortex (PCC), medial prefrontal cortex (mPFC)) are much less frequently implicated (i.e., only in 50% of the focused attention meditation literature) (Ganesan et al., 2022a). This drop in consistency across studies pertaining to brain regions compared to brain networks could be partially attributed to signal quality limitations inherent in 3 Tesla fMRI.

The advent of high strength 7 Tesla fMRI has the potential to ascertain group-level fMRI BOLD effects that are more reliable and neuroanatomically precise at the level of brain regions, compared to its lower strength 3 Tesla counterpart. This is because 7 Tesla fMRI enables MRI acquisition with higher neuroanatomical resolution, and stronger signal quality (i.e., 3 times higher signal-to-noise ratio) compared to 3 Tesla fMRI (Beisteiner et al., 2011; Gizewski et al., 2007; Hale et al., 2010; Pohmann et al., 2016; Theysohn et al., 2013; Trattnig et al., 2018). Emerging evidence also suggests that task-based fMRI data acquired with 7 Tesla can afford greater statistical power via producing reliable group-level results with fewer participants than 3 Tesla fMRI (Torrisi et al., 2018; Viessmann & Polimeni, 2021). Despite the technical advantages of 7 Tesla over 3 Tesla fMRI, there are no published 7 Tesla neuroimaging studies investigating meditation thus far to our knowledge.

Another issue in the emerging fMRI literature on focused attention meditation is the lack of adequate consideration of factors that may additionally affect fMRI responses measured during meditation. Specifically, most studies do not account for how physiological artifacts (e.g., cardiac and respiratory activity) affect fMRI findings (see review by Ganesan et al. (2022a) for details). This is particularly important because meditation (including focused attention meditation) is entrenched with physiological responses such as lowered heart rate, deeper and slower breathing, and lowered blood pressure (Ahani et al., 2013; Delmonte, 1984; Ditto et al., 2006; Soni & Muniyandi, 2019). Furthermore, physiological response fluctuations during fMRI task conditions can induce non-neuronal BOLD fMRI changes that can be mistaken for actual neuronal responses (Birn et al., 2006; Birn et al., 2009; Ganesan et al., 2022b). Therefore, lack of physiological artifact removal or correction during fMRI analysis can impact conclusions about the neurobiological underpinnings of meditation, including focused attention meditation. Similarly, fMRI activity in some brain areas within networks such as Default-mode and Executive Control may additionally be influenced by other non-physiological sources, such as inter-individual variability in dispositional mindfulness (Dickenson et al., 2013; Doll et al., 2016; Mooneyham et al., 2017; Scheibner et al., 2017), and level of arousal and effort during meditation tasks among beginners (Britton et al., 2014). Therefore, controlling for these measures that are entrenched with brain responses to meditation can potentially enable separating the neurobiological underpinnings of meditation from other attributes that may influence meditation performance.

Finally, many fMRI studies of meditation (including focused attention meditation) do not necessarily include assessments to measure behavioural changes before and after a meditation session inside an MRI scanner (Engström et al., 2022). Similarly, there is no clear understanding of how the behavioural impact produced by meditation inside the MRI scanner changes over time. Consequently, this poses a challenge in ascertaining the real-world impact of meditating inside an MRI scanner, and whether participants, especially beginners, can follow the meditation instructions as expected in the scanner. Preliminary evidence shows that a single brief session of meditation outside the MRI scanner can have positive effects on mindfulness levels (Johnson et al., 2015; Kim et al., 2019; Mrazek et al., 2012), mood (Broderick, 2005; Johnson et al., 2015), habitual psychological patterns (Wenk-Sormaz, 2005), emotion regulation (Arch & Craske, 2006), stress (Mohan et al., 2011), working memory (Yamaya et al., 2021), and executive function (Müller et al., 2021). However, it is unclear how a single-session of meditation inside the MRI scanner affects behaviour over time outside the scanner. The primary aim of our pilot study was to investigate the feasibility of using ultra-high strength (7 Tesla) fMRI via replication of core neuronal findings pertaining to focused attention meditation, using a small sample of beginner meditators (N=10). Based on aforementioned emerging evidence from 3 Tesla fMRI on focused attention meditation (Fox et al., 2016; Ganesan et al., 2022a), we hypothesised that 7 Tesla fMRI would enable robust detection of significantly reduced activation in core Default-mode network regions (e.g., PCC, mPFC) during focused attention meditation relative to non-focused rest, beyond physiological responses, subjective arousal, subjective effort and dispositional mindfulness.

Our secondary aim was to measure the physiological differences between focused attention meditation and non-focused rest during fMRI acquisition. We hypothesised that focused attention meditation would be accompanied by significantly slower breathing rate and heart rate, signifying physiological relaxation. Our additional exploratory aim was to measure how a single session of focused attention meditation in the MRI scanner impacts mindfulness-related outcomes outside the scanner longitudinally for up to 2 weeks (i.e., pre-fMRI to post-fMRI changes in state mindfulness, capacity for sustained attention, state anxiety and mind-wandering).

## 2 Methodology

We recruited 10 volunteers who were beginner meditators (4 males, 6 females; age = 30.1 ± 10.6 years) and free from major medical and psychiatric disorders via email advertisements from the local community. All volunteers provided written informed consent to participate, and the study was approved by the University of Melbourne human research ethics committee (Ethics ID: 22083).

### 2.1 Sample inclusion and exclusion criteria

The inclusion criteria were: (i) age between 19 and 60 years; (ii) an interest in learning and practicing meditation; (iii) fluency in English; and (iv) beginner at meditation, defined as having a cumulative lifetime meditation experience under 50 hours, with maximum weekly practice of 40 minutes over the past 6 months. The exclusion criteria were: (i) any lifetime clinical diagnoses of neuropsychiatric (e.g., psychosis, addictions, depression, anxiety) or neurological (e.g., traumatic brain injury, epilepsy) disorders, (ii) lifetime consumption of any psychoactive medication (e.g., antidepressants, benzodiazepines, anti-psychotics); or (iii) endorsement of any contraindications to MRI scanning.

### 2.2 Study procedure

This study comprised two main parts – meditation inside the MRI scanner, and out-of-scanner assessments to measure longitudinal behavioural changes associated with the fMRI meditation session (see Fig. 1).

**Fig. 1.**
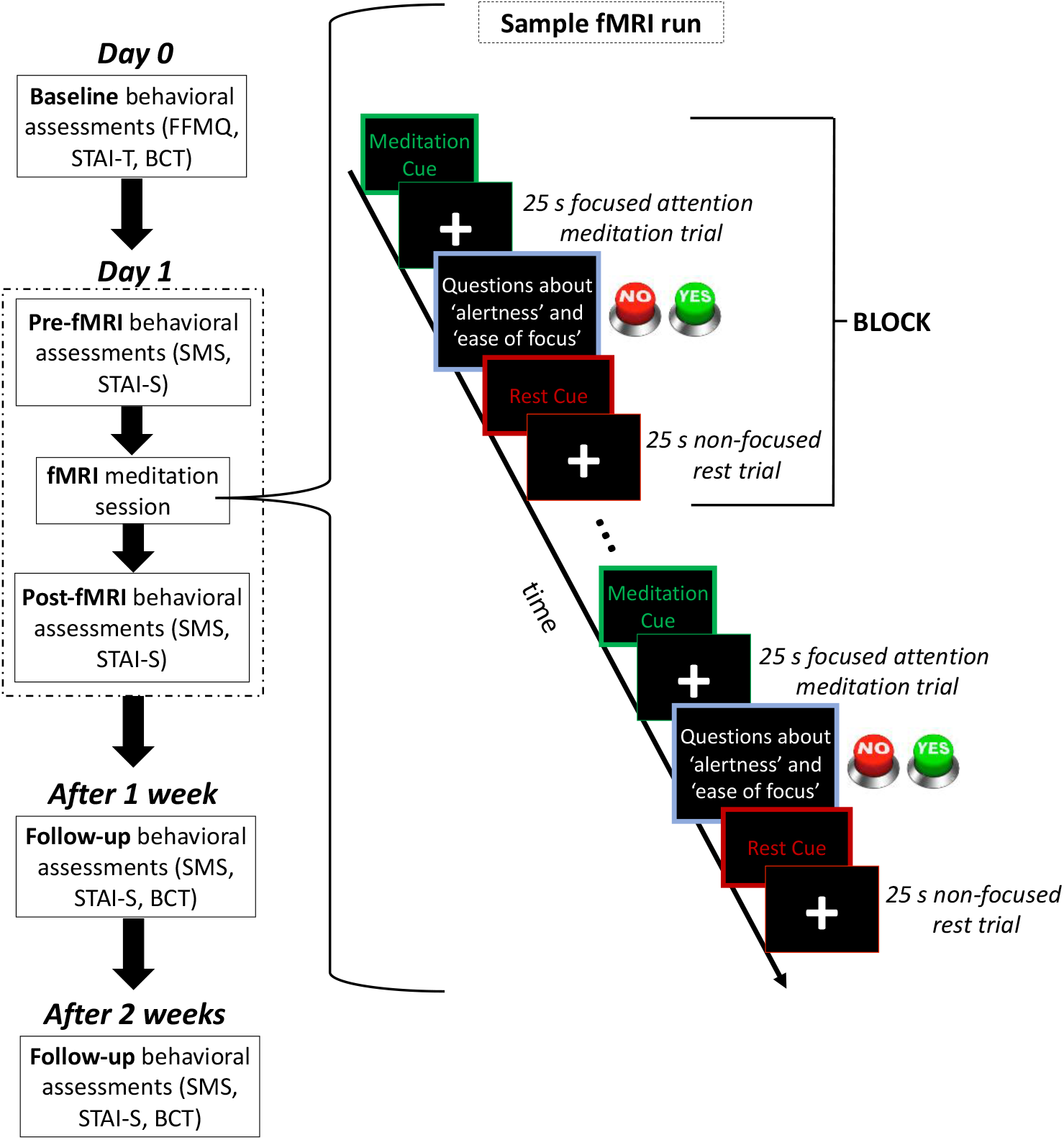
Graphical representation of the overall study paradigm along with a sample fMRI run. The overall study paradigm (as shown on the left) includes multiple measurement timepoints to assess longitudinal changes in behavioural measures outside the MRI scanner. FFMQ and STAI-T were only administered at baseline to characterize the recruited sample’s trait measures of mindfulness and anxiety. BCT with probes was administered 1 day before, 1 week after and 2 weeks after the fMRI meditation session. SMS and STAI-S were administered immediately before, immediately after, 1 week after and 2 weeks after the fMRI meditation session. Note that during each weekly follow-up, participants completed the SMS and STAI-S after completion of BCT with probes. The 7 Tesla fMRI meditation session consisted of 3 runs, where each run (as shown on the right) had 6 task blocks and 6 instances of button responses to ‘yes/no’ questions. Each task block comprised one non-focused rest trial of 25 s and one focused attention meditation trial of 25 s. FFMQ – Five Facet Mindfulness Questionnaire, STAI-T – State and Trait Anxiety Inventory - Trait module, BCT – Breath Counting Task, SMS – State Mindfulness Scale, STAI-S – State and Trait Anxiety Inventory - State module.

#### 2.2.1 MRI experimental design

Prior to MRI scanning, participants were familiarized with the focused attention meditation task and the control task in a mock scanner setup. For the fMRI scan, they were presented detailed instructions (adapted from Arch and Craske (2006)) about the two types of task conditions to be followed in an alternating order with their eyes open. Instructions for the focused attention meditation condition were: “*Focus on the actual sensations of breath entering and leaving the body. There is no need to think about the breath or change it. Just experience the sensations of it as you breathe in and out. When you notice that your awareness is no longer on the breath, gently bring your awareness back to the sensations of breathing*.” The instructions for the non-focused rest (non-meditation control) condition were: “*Lie still and simply think about whatever comes to mind, like usual throughout the day. Don’t focus on anything in particular*.”

Following the anatomical MRI scan, participants completed three fMRI runs, where each run comprised 6 task blocks (1 meditation trial and 1 rest trial in each task block) (see Fig. 1 for sample fMRI run). Each trial lasted for 25 seconds, and at the end of each meditation trial, participants were instructed to respond via buttons to two ‘Yes or No’ questions evaluating alertness and effort during the recent meditation trial (Q1. “*Was it easy to maintain your focus on the breath?*”; Q2. “*Did you feel quite sleepy/tired?*”). In-scanner alertness and in-scanner effort scores for each participant were calculated by averaging the number of affirmative responses to each respective question across trials and runs.

This study involves beginner meditators who are prone to mind-wandering distraction and attentional instability (Lomas et al., 2015; Lutz et al., 2008). Therefore, the short duration of trials (25 s) potentially minimized the occurrence of rest-like distraction during focused attention meditation conditions. Each task fMRI run lasted for approximately 10 minutes. Participants were scanned inside the MRI scanner for about 1 hour in total.

#### 2.2.1 MRI data acquisition

All MRI data was acquired on a 7 Tesla MRI scanner (Siemens Magnetom 7T plus) at the Melbourne Brain Centre Imaging Unit (MBCIU) using an 8/32 PTX/RX channel head coil, while timed visual display of cues and instructions inside the scanner was presented using the MATLAB Psychtoolbox software (version 3.1). A high resolution, RF inhomogeneity corrected and denoised (O’Brien et al., 2014) T1-weighted anatomical image (3D-MP2RAGE; 0.75mm x 0.75mm x 0.75mm; TE/TR = 2ms/5000ms) was acquired for post-hoc spatial registration with functional images. Functional images covering the whole brain were acquired using a multiband gradient-echo EPI sequence (Moeller et al., 2010) (1.6mm x 1.6mm x 1.6mm; TE/TR = 22ms/800ms; multiband acceleration = 6; field-of-view = 208 mm; matrix size = 130 × 130; 84 slices; slice thickness = 1.6mm; flip angle = 45^0^ ; P-A phase encoded). Concurrent to fMRI acquisition, respiratory signals of participants were recorded using a Siemens MRI compatible respiration belt worn around the abdomen, and cardiac measurements were recorded using a Siemens MRI compatible pulse oximetry sensor worn on a fingertip.

#### 2.2.1 Self-report behavioural assessments

At baseline, we characterised the sample by age, sex and self-reported lifetime meditation experience (in hours), dispositional mindfulness and trait anxiety. Dispositional mindfulness was measured only at baseline by administering the Five Facet Mindfulness Questionnaire (FFMQ; Baer et al., 2006). The FFMQ comprises 39 self-report questions covering 5 mindfulness facets, i.e., observing, describing, acting with awareness, non-reactivity to inner experiences, and non-judging of inner experiences (Baer et al., 2006). This questionnaire is scored out of 5, with higher scores suggesting greater dispositional mindfulness. An overall FFMQ score from averaging across all sub-scales represents the overall multi-faceted mindfulness trait of an individual. Trait anxiety was measured only at baseline with the State and Trait Anxiety Inventory, Trait module (STAI-T). The STAI-T comprises 20 rating-based questions, and measured trait levels of anxiety in each participant (Spielberger et al., 1983). This questionnaire is scored out of 4, with lower scores indicating lower trait levels of anxiety. We measured changes in self-reported state mindfulness and state anxiety pre- to post-fMRI meditation for up to 2 weeks. Specifically, these measures were administered immediately before and after the fMRI meditation session, as well as 1 week and 2 weeks after the fMRI session (Fig. 1). State mindfulness was measured with the State Mindfulness Scale (SMS) (Tanay & Bernstein, 2013), which (Spielberger et al., 1983) comprises 21 statements with 5-point ratings to measure state mindfulness. An average SMS score of 5 represents the most mindful state. State anxiety was measured with the state anxiety subscale from the STAI.

STAI-S has 20 statements with 4-point ratings assessing state anxiety, where an average rating score of 4 represents the most stressful state.

#### 2.2.1 Computerized behavioural assessment

We objectively measured participants’ ability to focus and sustain attention on the breath at baseline before fMRI meditation, and 1 week and 2 weeks after fMRI meditation (Fig. 1). This was measured using a standardized 20-minute computerized breath counting task (BCT) (Levinson et al., 2014) with experiential probes (Frewen et al., 2008). Due to good test-retest reliability for a 1-week interval (Levinson et al., 2014), we required participants to perform this task three times with a gap of at least one week during the course of the study.

In this task, participants were instructed to count their breaths from 1 to 9 cyclically with their eyes closed. The first 8 counts were accompanied by left arrow key presses, while the 9^th^ and last breath of each BCT cycle was indicated by a right arrow key press. Instances of self-caught miscounting due to mind-wandering distraction were accompanied by ‘Shift’ key presses, which restarted the BCT cycle from 1. At six pseudo-random instances during BCT, participants were probed to verbally report their most recent breath count number, and to check if their attention was focused on the breath at that moment (based on verbal yes/no response to ‘Was your attention on the breath just now?’). The physiological veracity of self-reported breath counts was evaluated via concurrent respiratory measurements using a commercial wearable respiration belt (Vernier Science Education, Oregon, USA).

The outcome measures of this task include: i) BCT accuracy (% of correct count cycles), ii) BCT miscount (% of incorrect count cycles), iii) BCT reset (% of reset count cycles), iv) BCT probe accuracy (% of affirmative probe responses). Higher BCT and probe accuracies, and lower BCT miscount percentage indicate better task performance due to reduced attentional lapses and less frequent mind-wandering distraction. Higher BCT reset percentage suggests greater meta-awareness of mind-wandering distraction, while also being potentially indicative of more frequent distraction (K et al., 2018).

### Analysis details

#### 2.3.1 FMRI data pre-processing

MRI images were acquired in the DICOM format and converted to the NifTI format using the dcm2niix tool (Li et al., 2016). Distortions in the fMRI EPI images due to magnetic field inhomogeneities were corrected using acquired reverse-phase encoded (A-P) EPI images with FSL topup (Andersson et al., 2003). Skull stripping and brain image extraction from the anatomical scans were performed using ANTs (Avants et al., 2009). Using Motion Correction FMRIB’s Linear Image Registration Tool (MCFLIRT) in FSL (Jenkinson et al., 2002), linear rigid-body transformation (rotation and translation) was performed on the fMRI images to correct for head motion. Each participant’s low-resolution motion-corrected fMRI images were then linearly co-registered to their respective high-resolution anatomical brain image (output from ANTs), and subsequently to the Montreal Neurological Institute (MNI) standard stereotactic space using FLIRT in FSL (Jenkinson et al., 2002; Jenkinson & Smith, 2001). Finally, using FSL FEAT (Woolrich et al., 2001), pre-whitening was applied to the voxel-wise fMRI BOLD timeseries to correct for temporal autocorrelation; temporal high-pass filtering (0.01 Hz) was used to remove low frequency noise; and spatial smoothing was applied using a Gaussian kernel size of 3.2 mm full-width half maximum (FWHM). A fast TR sampling of 800 ms with multiband acceleration precluded the need for slice-timing correction (M. F. Glasser et al., 2013).

#### 2.3.2 Functional brain activation during focused attention meditation relative to non-focused rest

Whole-brain analyses were undertaken through GLM to identify brain areas that significantly increased or decreased activation during focused attention meditation relative to non-focused rest. This was performed using FSL FEAT (Woolrich et al., 2001) and Permutation Analysis of Linear Models (PALM) (Winkler et al., 2014).

The time course of each condition (meditation, rest, cue/instructions, and button responses as shown in Fig. 1 sample fMRI run) was convolved with the canonical double-gamma hemodynamic response function (HRF), temporally smoothed (0.01 Hz; same as data), and entered as a block design predictor to model the voxel-wise fMRI BOLD timeseries of each run separately. The temporal derivatives of each of these four condition predictors were also included to improve overall model fit. Additionally, head motion artifacts in every voxel’s blood oxygen level-dependent (BOLD) timeseries were accounted for by including six motion parameters (3 rotation and 3 translation; from MCFLIRT) and their respective derivatives as nuisance predictors in the model. Nuisance predictors modelling large and sudden motion (generated by FSL Motion Outliers tool (https://fsl.fmrib.ox.ac.uk/fsl/fslwiki/FSLMotionOutliers)) which affect specific BOLD timepoints were also included in the model.

To correct for physiological artefacts during first-level GLM, RETROspective Image CORrection (RETROICOR) (Glover et al., 2000) was performed on the BOLD timeseries using the PhysIO toolbox with the respiration and cardiac data acquired during fMRI scanning. Specifically, 22 physiological nuisance predictors, accounting for respiration signals (8 regressors), cardiac signals (6 regressors), interaction between respiration and cardiac signals (4 regressors), heart rate (1 regressor) (Chang et al., 2009), and respiratory volume per time and their time-shifted values (3 regressors) (Harrison et al., 2021), were included in the whole-brain voxel-wise GLM of each fMRI run from each participant. Parameter estimates for the linear model fit were calculated for the contrast of meditation relative to. rest.

Subsequently, outputs from the first-level analysis were entered into second-level GLM in FEAT to calculate the average voxel-wise response across runs for each participant. For group-level inferences, outputs from the second-level GLM were further entered into an across-participant third-level GLM in PALM. As covariates, participant-level average measures of self-reported alertness score and effort score during the meditation task (from the in-scanner button responses), as well as overall baseline dispositional mindfulness (total baseline FFMQ score) were included in the group-level GLM (see Supplementary Fig. S3-S5 for GLM design matrices). These covariates were included at the group-level to control for the influence of inter-individual variability in arousal and effort during the meditation task, as well as general baseline trait mindfulness (FFMQ). Following third-level GLM, clusters of spatially contiguous voxels were delineated after thresholding the voxels at z=3.1 (uncorrected cluster-forming p<0.001). Statistically significant clusters of activation and deactivation during meditation (compared to non-task rest) were determined through accelerated non-parametric permutation testing (1024 permutations of sign-flips) (Nichols & Holmes, 2002; Winkler et al., 2016), along with family-wise error (FWE) control for multiple comparisons across clusters (Alberton et al., 2020) at p<0.05. For GLM group-level results without the inclusion of abovementioned covariates (i.e., arousal, effort and dispositional mindfulness), refer to Supplementary Fig. S1.

#### 2.3.3 Physiological differences during 7T fMRI between focused attention meditation and non-focused rest

Repeated-measures general linear modelling (GLM) was implemented in MATLAB to test for statistically significant differences in key respiration and cardiac measures recorded during the 7 Tesla fMRI session between focused attention meditation and non-focused rest conditions. Specifically, the PhysIO toolbox (Kasper et al., 2017) was used to calculate respiration rate, respiratory volume per time (Harrison et al., 2021), and heart rate (Chang et al., 2009) values for each BOLD timepoint based on the physiological data acquired during 7 Tesla fMRI scanning. Subsequently, for each physiological measure, a mean trial-wise value was calculated by averaging across the physiological values within each condition trial (meditation or rest). For every participant and each measure, this produced 6 mean trial-wise values for meditation and 6 for rest within each run. 3 independent one-way repeated-measures Analyses of Variance (ANOVA) were used to test for differences in each respective physiological measure (i.e., respiration rate, respiratory volume per time, and heart rate) between meditation and rest. Specifically, for each ANOVA, mean trial-wise values from every run and participant were used. Statistical significance of differences was assessed after Bonferroni control for multiple comparisons (p<0.017).

#### 2.3.4 Longitudinal changes in behavioral assessments following fMRI meditation

Longitudinal pre- to post-fMRI changes in state mindfulness, state anxiety and outcome measures of BCT were quantified through independent one-way repeated measures ANOVA (with time as independent variable) and Bonferroni correction for multiple comparisons across analyses (p < 0.0083). Specifically, 2 independent ANOVA were used to examine group-level changes from baseline to follow-ups in SMS and STAI-S scores measured at 4 time points, i.e., pre-fMRI, post-fMRI, 1-week follow up, and 2-week follow up. Similarly, 4 independent ANOVA analyses were used to examine group-level changes in breath attention probe accuracy, BCT accuracy, BCT resets and BCT miscounts across 3 time points, i.e., pre-fMRI, 1-week follow up and 2-week follow up.

## 3 Results

### 3.1 Sample characteristics at baseline

The recruited sample primarily comprised young adults with cumulative lifetime meditation experience ranging between 0 and 50 hours. Based on average and median scores, the included beginner meditators sample (N=10) had low levels of trait anxiety and moderate levels of dispositional mindfulness. The mean, standard deviation, median and range of key sample demographics, anxiety and mindfulness levels are listed in Table 1.

**Table 1:** Key characteristics of the recruited sample (N=10), reported as mean, standard deviation, median and range.

| Measure | Sample mean ± standard deviation<br>(median [range]) |
| --- | --- |
| Age | 30 ± 11 (27 [21-57]) years |
| Trait anxiety STAI-T score | 1.9 ± 0.5 (1.8 [1.4-3.2]) |
| Self-reported lifetime meditation experience | 20.6 ± 22.5 (11.5 [0-50]) hours |
| Trait mindfulness – FFMQ observing score | 3.3 ± 0.6 (3.4 [1.9-4.0]) |
| FFMQ describing score | 3.4 ± 0.7 (3.5 [2.4-4.5]) |
| FFMQ acting with awareness score | 3.2 ± 0.7 (3.5 [2.0-4.2]) |
| FFMQ non-judging score | 3.8 ± 0.8 (3.8 [1.7-4.7]) |
| FFMQ non-reactivity score | 2.9 ± 0.4 (3.1 [2.3-3.6]) |
| FFMQ overall score | 3.3 ± 0.5 (3.6 [2.2-3.8]) |

*FFMQ – Five Facet Mindfulness Questionnaire; STAI-T – State and Trait Anxiety Inventory - Trait module*

### 3.2 fMRI brain activation during focused attention meditation relative to non-focused rest

After FWE correction across clusters, the whole-brain GLM analyses revealed several significant group-level deactivation clusters during focused attention meditation compared to rest condition (non-meditation control). These significant clusters comprised posterior insula, anterior cingulate cortex (ACC), hippocampal areas, cerebellum, posterior cingulate cortex, medial prefrontal cortex, precuneus, visual cortex and thalamus (see Supplementary Fig. S1 and explanation for details).

After further control for inter-individual variability in baseline dispositional mindfulness (total baseline FFMQ score), and average in-scanner alertness and average in-scanner effort during meditation, significant deactivation clusters were confined to the occipital cortex, thalamus (lateral ventral/dorsal posterior nuclei) and Default-mode network, i.e., precuneus, posterior cingulate cortex and antero-medial prefrontal cortex (shown in Fig. 2 and Table 2).

**Fig. 2.**
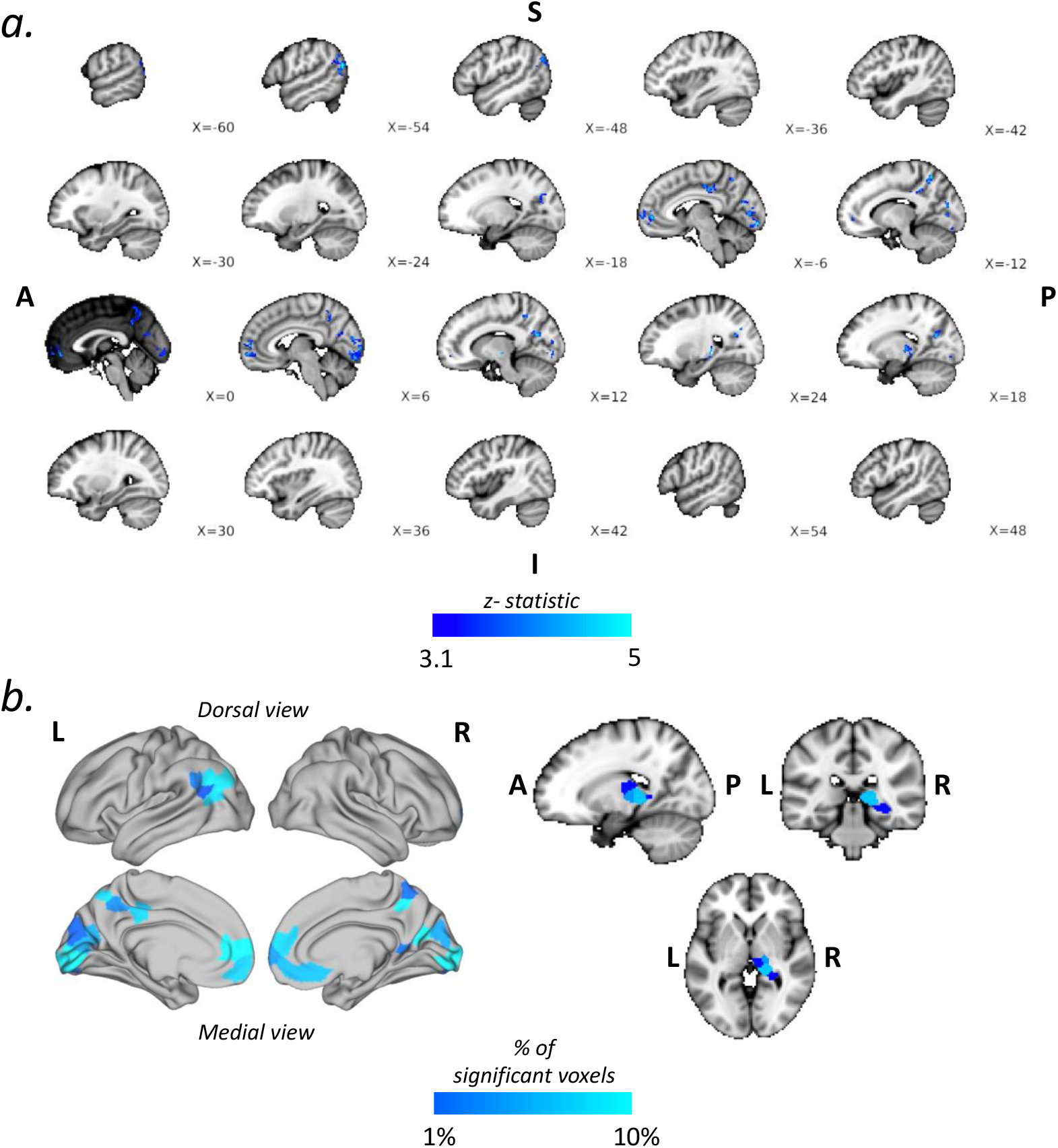
Representation of brain clusters showing significant functional deactivation during MEDITATION relative to REST with general linear modelling (GLM) analysis, after controlling for overall baseline trait mindfulness, in-scanner alertness and in-scanner effort during meditation. **a)** Section of volumetric sagittal brain slices along with x-coordinates displaying the significantly deactivated brain areas during meditation relative to rest. The z-statistic value corresponding to the magnitude of deactivation determines a region’s colour (**‘cool’ colour gradient**). **b)** Brain display showing the percentage of voxels from the standard 400-region Schaeffer-Tian template that overlapped with the significant deactivation clusters during MEDITATION relative to REST. The left panel shows cortical overlap percentages mapped on to the brain surface, while the right panel shows subcortical overlap percentages via discrete anatomical slices. The brain surface mapping was performed with the Glasser brain surface template (Glasser et al., 2016). The percentage of overlapping significant voxels determines a region’s colour (**‘cool’ colour gradient**). S – superior/dorsal, A – anterior, P – posterior, I – inferior/ventral.

**Table 2:** Overview of brain regions in standard MNI space showing significantly reduced activity during MEDITATION relative to REST, controlling for total baseline FFMQ score, average in-scanner alertness score and average in-scanner effort score.

| Significantly deactivated areas during Meditation relative to Rest | Cluster size (voxels) | Cluster peak value (z-stat) | Cluster significance (FWE p-value) | Cluster peak location (MNI) |  |  |
| --- | --- | --- | --- | --- | --- | --- |
|  |  |  |  | X | Y | Z |
| bilateral occipital cortex - lingual gyri, fusiform gyri, calcarine cortex, occipital pole | 442 | 4.4 | 0.0007 | -8 | -86 | -12 |
| medial prefrontal cortex, paracingulate gyrus, frontal pole | 302 | 4.5 | 0.0046 | -4 | 52 | 0 |
| R. precuneus, R. cuneus | 270 | 4.4 | 0.0071 | 14 | -66 | 22 |
| L. precuneus, posterior cingulate gyrus | 223 | 4.2 | 0.0129 | -12 | -58 | 56 |
| L. lateral occipital complex, L. angular gyrus | 207 | 4.9 | 0.0159 | -54 | -70 | 22 |
| R. lateral ventro/dorsoposterior thalamus | 121 | 4.6 | 0.0470 | 22 | -30 | 0 |
| L. posterior cingulate gyrus | 121 | 4.3 | 0.0470 | -8 | -28 | 38 |

L – left, R – right

On the other hand, there were no significant activation clusters during focused attention meditation relative to rest, before and after controlling for overall FFMQ, alertness and effort. Furthermore, there were no significant correlations between the significant deactivation clusters and total FFMQ score, in-scanner alertness score during meditation or in-scanner effort score during meditation.

### 3.2 In-scanner physiological differences between focused attention meditation and non-focused rest

As hypothesised, the respiration rate (breaths per minute), respiratory volume (volume per minute) and heart rate (beats per minute) were significantly different (after Bonferroni correction, p<0.017) between focused attention meditation and non-focused rest during the 7 Tesla fMRI session across participants (Fig. 3).

**Fig. 3.**
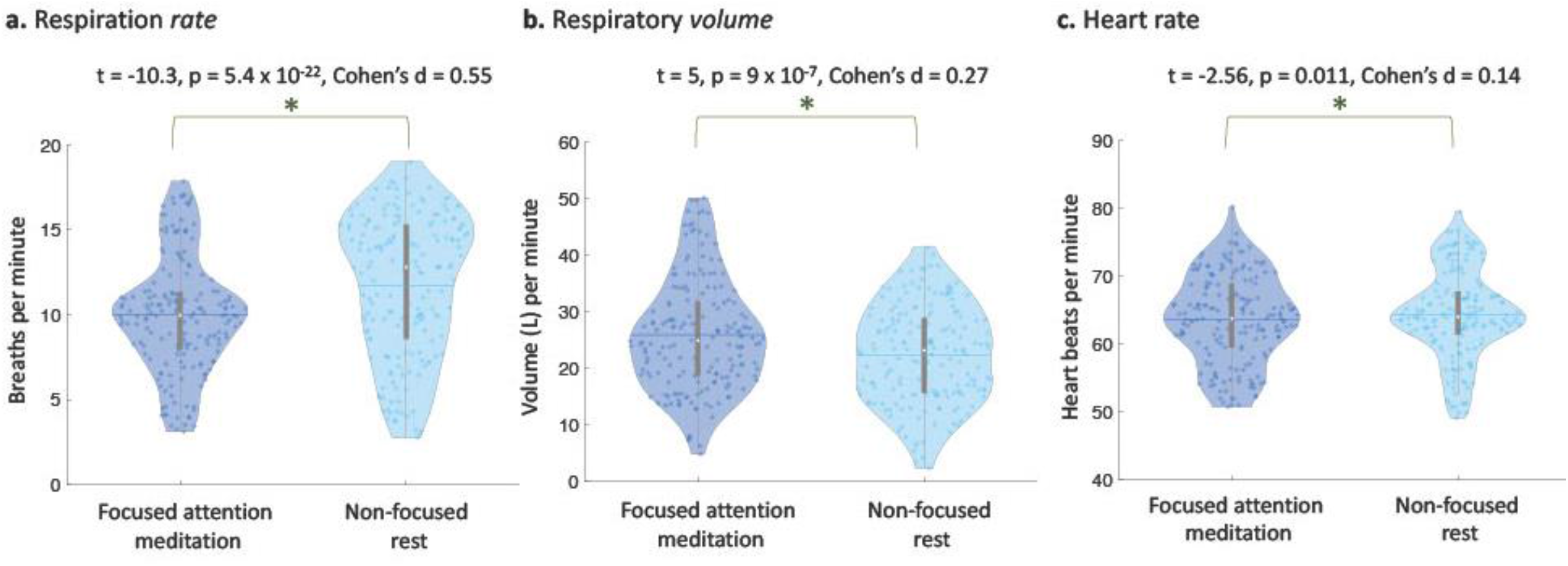
Violin plots comparing distributions of respiration rate, respiratory volume and heart rate from each fMRI task condition, i.e., focused attention meditation and non-focused rest. All comparisons were performed with one-way repeated-measures ANOVA. **a)** Respiration rate for each trial within each task condition (meditation and rest) from every participant. Each coloured data point within a violin plot of a specific condition represents the average respiration rate during a corresponding trial from a participant. The median value of each plot is indicated by a white dot at the centre. The mean is indicated by the horizontal line passing through the thickness of the plot. We found that the respiration rate was significantly lower during meditation relative to rest across trials and participants. **b)** Same as (a) but for respiratory volume. We found that the respiratory volume inhaled was significantly higher during meditation relative to rest across trials and participants. **c)** Same as (a) but for heart rate. We found that the heart rate was significantly lower during meditation relative to rest across trials and participants.

The maximum difference was observed in the respiration rate, which was significantly lower during meditation relative to rest (t = -10.3, p = 5.4 × 10^−22^, Cohen’s d = 0.55). On the other hand, the volume of air breathed (respiratory volume) was significantly greater (t = 5, p = 9 × 10^−7^, Cohen’s d = 0.27) during meditation. Similarly, albeit by a smaller magnitude, the heart rate also significantly decreased during meditation relative to rest (t = -2.56, p = 0.011, Cohen’s d = 0.14). Note that these physiological differences have been accounted for in all fMRI analyses in this study. In Supplementary Fig. S2, we further demonstrate the effect of physiological confounds by comparing the extent of significant brain de-activation areas found here with those obtained *without* physiological artifact correction.

### 3.4 Longitudinal pre- to post-fMRI change in outside scanner behavioural measures up to 2 weeks

Only state mindfulness measured by SMS showed a significant longitudinal improvement (F = 8.16, p = 0.0005; survived Bonferroni correction) following the 7 Tesla fMRI meditation session compared to baseline. Specifically, compared to baseline, there was a significant rise in state mindfulness across participants immediately following the single 7 Tesla fMRI meditation session, with effects maintained for up to 2 weeks after the session (Fig. 4a).

**Fig. 4.**
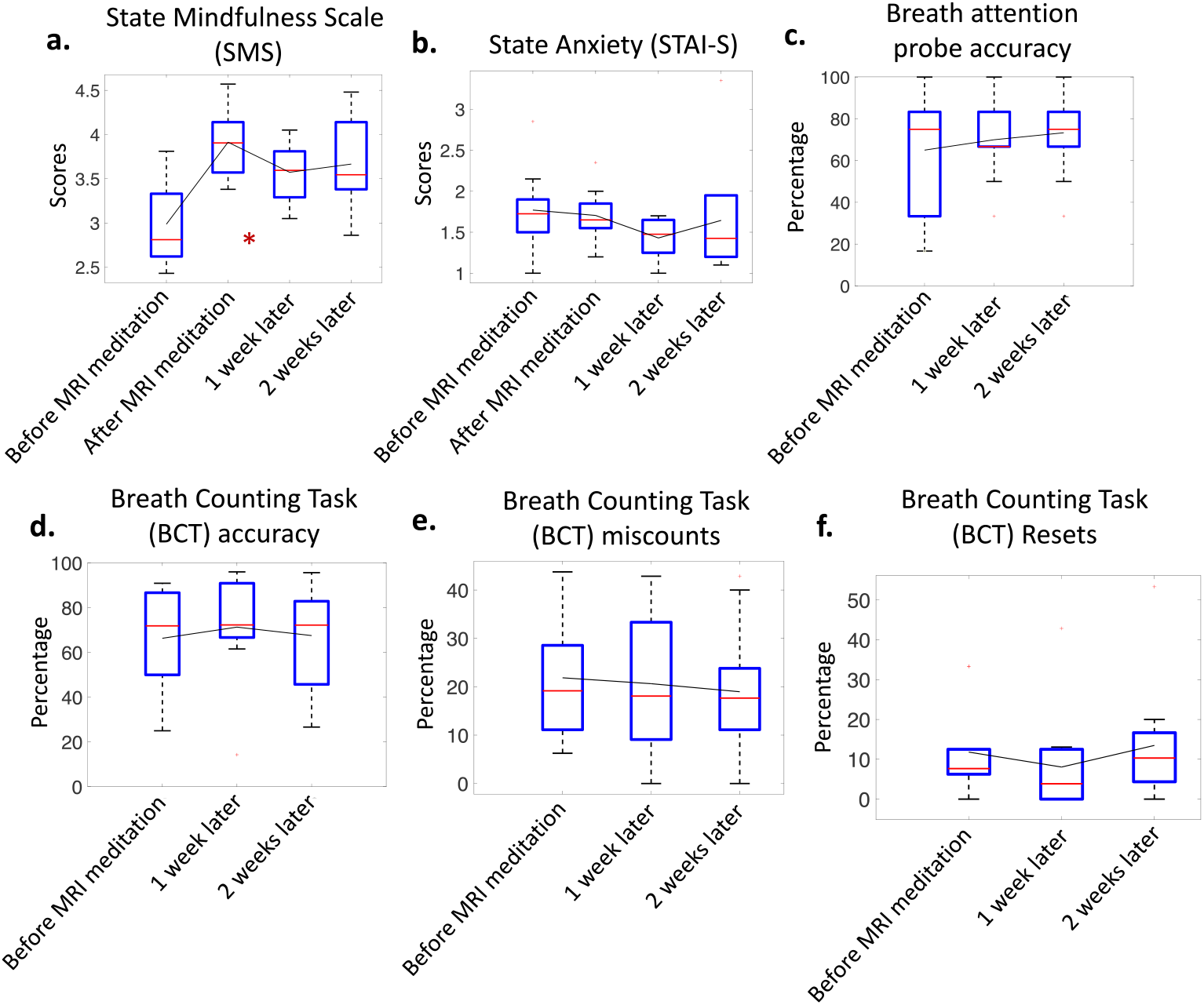
Bar graphs representing group-level longitudinal changes in behavioural measures following a single session of 7 Tesla fMRI meditation. a) Changes in SMS scores across N=10 participants over 4 time points, i.e., immediately before (baseline), immediately after, 1 week after and 2 weeks after fMRI meditation session (significant after Bonferroni correction (p < 0.05/6). b) Same as (a) for STAI-S scores and the changes were non-significant. c) Changes in breath attention probe accuracy percentages across N=10 participants over 3 time points, i.e., before (baseline), 1 week after and 2 weeks after fMRI meditation session. The changes were not significant. d) Same as (c) for BCT accuracy percentages. e) Same as (c) for BCT miscount percentages. f) Same as (c) for BCT reset percentages. In each bar graph, the bars corresponding to different time points are joined together at their respective sample means. The sample median is indicated by the red line within each bar. The extended dotted lines in each bar represent the 75% confidence interval. * - significant trend after Bonferroni correction.

The other measures showed some non-significant changes over time. Compared to baseline, average STAI-S scores (F = 1.44, p = 0.25) showed a slight decrease 1 week after the fMRI meditation session, which was not maintained at the 2-week mark (Fig. 4b). The breath attention probe accuracy (F = 0.66, p = 0.53) also showed non-significant increasing trend on average, with diminishing across-participant variability in accuracies following fMRI meditation compared to baseline (Fig. 4c). During each BCT session, breathing rate was found to be significantly and strongly correlated with breath count rate (r1 = 0.97, p = 1.5×10^−6^; r2 = 0.93, p2 = 8.2×10^−5^; r3 = 0.99, p3 = 1.1×10^−7^), thus physiologically verifying the validity of breath counts. Participants showed a slight average improvement in BCT accuracy after 1 week with a return to baseline at the 2-week mark (F = 0.42, p = 0.66; Fig. 4d). Compared to baseline, the average BCT reset percentages showed a decline after 1 week with a small rise again after 2 weeks following fMRI meditation (F = 1.96, p = 0.169; Fig. 4f). On the other hand, BCT miscount percentages showed negligible group-level longitudinal change (F = 0.16, p = 0.85; Fig. 4e).

## 4 Discussion

The current pilot study is one of the first to examine the neural substrates of meditation using ultra-high strength 7 Tesla functional MRI. Specifically, we examined the feasibility of investigating focused attention meditation with a small sample of beginner meditators (N=10) scanned using high power 7 Tesla fMRI. After controlling for physiological responses, head motion, baseline dispositional mindfulness, subjective arousal during meditation and subjective effort during meditation, we found significantly decreased activation (de-activation relative to rest) of Default-mode network regions (mPFC, precuneus, PCC) during meditation relative to rest. These Default-mode deactivations likely suggest attenuation of mind-wandering and spontaneous thought during meditation. Our pilot 7 Tesla fMRI findings hence successfully replicate existing widespread evidence implicating deactivation of the Default-mode network during focused attention meditation, despite the inclusion of conservative statistical corrections in a small sample. We also found significant deactivation in thalamic nuclei and occipital cortex during meditation relative to rest suggesting a role in perceptual decoupling during meditation. On the other hand, there were no brain areas that showed increases in activation during meditation relative to rest. Among measured behavioural attributes, compared to baseline, we observed a significant elevation in state mindfulness following the fMRI meditation session and this level was closely maintained at subsequent weekly follow-ups for 2 weeks. Although we observed no significant changes from baseline in state anxiety and focused attention (breath counting task outcomes), these attributes demonstrated slight non-linear trends over time following the fMRI meditation session.

### 4.1 Default-mode network areas are attenuated during focused attention meditation

We found that focused attention meditation deactivates circumscribed areas within PCC, precuneus, and anterior mPFC, i.e., key nodes of the Default-mode network, relative to non-focused resting-state. These deactivations were significant after accounting for overall dispositional mindfulness (overall baseline FFMQ score), self-reported effort during fMRI meditation and self-reported alertness during fMRI meditation (average affirmative responses after meditation trials). When these measures were not included as covariates in the fMRI analyses, significant deactivations during focused attention meditation relative to rest were more diffuse and less specific (See Supplementary Fig. S1 and explanation for more details). Thus, it is possible that the more specific Default-mode deactivations reported herein are not influenced by inter-individual differences in overall mindfulness ability, and momentary levels of arousal and effort during the meditation task.

Our findings suggest that deactivations of key Default mode network regions relative to rest may diminish brain activity implicated in self-referential processing, mental predictions, repetitive thought and mental time-travel during focused attention meditation. Activity in Default-mode network regions has been widely associated with mind-wandering and spontaneous thought (Fox et al., 2015). These regions typically show the highest metabolic activity at baseline, and their metabolic activity diminishes with goal-directed cognition or perception (Raichle et al., 2001). Notably, focused attention meditation also includes goal-directed perception as it involves directing attention away from mind-wandering and cognition towards a target object (e.g., breathing sensations) or experience in the present moment. Therefore, goal-directed perception of the target object/experience during focused attention meditation may have been facilitated by diminishing activation of regions within the Default-mode network relative to non-focused resting-state, i.e., PCC, precuneus and mPFC (Fox et al., 2016; Ganesan et al., 2022a). The PCC is most commonly implicated in self-directed cognition, spontaneous thought and conceptual processing (Leech & Sharp, 2014; Leech & Smallwood, 2019). Similarly, the precuneus is a hub for memory processing and mental representations of an integrative self, and shows hypoactivity during sleep, hypnosis and sedation (Cavanna & Trimble, 2006; Utevsky et al., 2014). Further, anterior mPFC is posited to underlie self-referential, value and reward processing and planning (Andrews-Hanna et al., 2010; Lieberman et al., 2019). Consequently, deactivations in these regions may free up attentional resources in order to improve quality of deliberate focus on present moment objects or experiences (Ganesan et al., 2022a; Laukkonen & Slagter, 2021).

Overall, our pilot 7 Tesla fMRI findings successfully replicate Default-mode deactivation relative to rest. This highlights the accessibility of focused attention meditation regardless of prior meditation experience, as well as the high sensitivity of 7 Tesla fMRI in capturing such core neurobiological underpinnings despite a small sample size and stringent statistical corrections.

### 4.2 Role of thalamic and occipital de-activations during focused attention meditation relative to rest

In addition to Default-mode network areas, we also found significant deactivations in right lateral posterior thalamic nuclei, and several occipital areas during focused attention meditation relative to non-focused rest. While thalamic subdivisions have shown increases in activation relative to rest during focused attention meditation (on breath) (Farb et al., 2013; Hasenkamp et al., 2012), decreases in thalamic activation have also been reported (Farb et al., 2013; May et al., 2016). Overall, the role of thalamus in the context of focused attention meditation is unclear. The thalamus is known to integrate and relay multimodal sensory information back and forth between the cortex and nervous system, thereby facilitating perception (Hwang et al., 2017). Particularly, first-order ventral/dorsal lateral posterior nuclei of the thalamus are involved in communicating somatosensory information (e.g., proprioception, touch and pain sensations) with the primary somatomotor cortex (Prescott & Ratté, 2017). During focused attention meditation, the perception and integration of somatosensory information and other stimuli is likely attenuated due to attention being exclusively directed towards a specific target (e.g., breathing sensations) (Laukkonen & Slagter, 2021). Such attenuation of non-target stimuli and sensations likely explains the circumscribed deactivations observed in the lateral posterior thalamic nuclei during focused attention meditation relative to rest in the current study.

We also found that several occipital regions were deactivated during focused attention meditation relative to rest. Each of these occipital areas deactivated in the current study have been widely implicated in visual recognition, processing and perception, i.e., lateral occipital complex (Grill-Spector et al., 2001), fusiform gyri (Weiner & Zilles, 2016), angular gyri (Seghier, 2013), occipital pole, calcarine area and lingual gyri (Swenson & Gulledge, 2017). Deactivations in the visual occipital cortex observed herein are consistent with majority of the extant fMRI literature on focused attention meditation (Baron Short et al., 2010; Dickenson et al., 2013; Farb et al., 2013; Hasenkamp et al., 2012; Hölzel et al., 2007; Ritskes et al., 2004; Scheibner et al., 2017). Occipital deactivations likely result from perceptual decoupling of non-target visual stimuli (e.g., fixation cross) presented during focused attention meditation fMRI trials, as participants likely tried to focus their attention on breathing sensations (target stimulus) while keeping their eyes open as instructed.

Note that we did not find any significant activations during meditation relative to rest within other brain regions typically expected to be involved in focused attention meditation, i.e., Salience and Executive Control network regions. The short duration of focused attention meditation trials may have minimized the scope for mind-wandering distraction. This likely minimized the need for Control network facilitated attentional switches between distraction and breath during meditation trials (Ganesan et al., 2022a). As a potential consequence, we did not find any significant group-level activations within the Control network during meditation relative to rest with our current sample. Similarly, the short meditation duration may have also been insufficient for beginners to distinctly perceive subtle breathing sensations. Hence, Salience network activations during meditation relative to rest in our current sample were possibly quite variable across participants and hence insignificant at the group level. Future study designs may benefit from inclusion of both short as well as longer meditation trials to additionally delineate the effects of meditation duration on distraction, attention and awareness.

### 4.3 Cardiac and respiratory changes during focused attention meditation

During focused attention meditation relative to non-focused rest, we found significant decreases in heart rate and breathing rate with a significant increase in volume of respiration as expected. This is consistent with previous reports of lowered physiological arousal during meditative compared to non-meditative control states, i.e., lowered heart rate and deepened breathing at a reduced pace (Ahani et al., 2013; Delmonte, 1984; Farb et al., 2013; Manna et al., 2010; Soni & Muniyandi, 2019; Weng et al., 2020).

On the other hand, such physiological response fluctuations between meditation and rest are considered artifacts in an fMRI context with the potential to alter significant findings (Birn et al., 2006; Birn et al., 2009; Ganesan et al., 2022b). Specifically, physiological measures that covary with task conditions (i.e., meditation and rest conditions here) can often conflate the source of fMRI BOLD responses. Consequently, some of the significant brain activations and deactivations can be missed (false-negatives) or misattributed (false-positives) to neuronally-induced task responses (Birn et al., 2009), which is potentially prevalent in most extant fMRI studies investigating focused attention meditation (Ganesan et al., 2022a).

Statistical corrections (RETROICOR regression) that removed linear effects of measured physiological signals, i.e., cardiac and respiratory signals, from fMRI BOLD responses, were included during fMRI analyses in this study. Inclusion vs. exclusion of physiological artifact correction demonstrates noticeable changes in the extent of de-activation during focused attention meditation relative to rest (see Supplementary Fig. 2). Specifically, notable differences can be observed in mid-line cortical areas that have been typically associated with focused attention meditation (such as PCC and precuneus). Our preliminary findings thus warrant the inclusion of physiological corrections in fMRI studies of focused attention meditation, to mitigate both false positive as well as false negative findings pertaining to fMRI brain function during meditation.

### 4.4 Longitudinal behavioural changes after a single fMRI meditation session: preliminary evidence

In our exploratory behavioural analyses, we observed a significant rise pre- to post-fMRI meditation in self-reported state mindfulness (SMS). This is potentially indicative of participants experiencing meditative states as expected during the fMRI session, thus leading to subsequent elevation in present-moment awareness of mental content and bodily sensations (Tanay & Bernstein, 2013). Increases in state mindfulness measured by SMS can predict improvements in dispositional mindfulness after several weeks (Tanay & Bernstein, 2013), which likely explains the elevated SMS scores for up to 2 weeks after the fMRI meditation session.

On the other hand, longitudinal changes in breath counting (BCT) and self-reported state anxiety (STAI-S) measures did not survive statistical significance. These measures however demonstrated slight non-significant trends of change over 2 weeks. Specifically, most of the breath counting task outcomes showed mild non-linear change compared to baseline, with increases after 1-week followed by decreases at the 2-week follow-up. The scope for significant decreases in state anxiety (STAI-S) may be limited in a small sample of normative individuals with low levels of baseline state and trait anxiety. Similarly, a single session of focused attention meditation in a small sample of beginner meditators may be insufficient to produce robust and significant impact on attention and mind-wandering (as measured by BCT).

### 4.5 Limitations

The findings from this 7 Tesla fMRI pilot study need to be interpreted in light of several methodological limitations. Firstly, the sample size used in this study is small (N=10). Although we replicated neural findings pertaining to focused attention meditation with this sample, adequately powered future studies are required to validate these findings and investigate subtler underlying neuronal responses. Furthermore, we did not find any direct significant associations between fMRI activity and overall baseline dispositional mindfulness, self-reported effort scores during meditation or self-reported arousal scores during meditation, likely due to the limited sample size. Similarly, we did not explore the associations between each facet of FFMQ and fMRI activity due to the small sample. Note that a larger sample for an adequately powered 7 Tesla fMRI study would ideally be lower compared to that for an equivalent 3 Tesla fMRI study (Torrisi et al., 2018; Viessmann & Polimeni, 2021).

Secondly, in addition to the small sample size, this study did not include a placebo control group for the behavioural measurements. Therefore, caution is required when interpretating the longitudinal behavioural effects observed in this pilot study. In the future, deeper understanding about the extent of longitudinal benefits conferred by a single meditation session using necessary control groups and adequate sample sizes could potentially minimise attrition from meditation practice especially among beginners and clinical populations.

Finally, our experimental design included a response block after meditation but not after rest condition. This could have led to differences in anticipation or outcome prediction between the two primary conditions (i.e., meditation and rest) being compared. However, these anticipatory effects may have been minimal, as brain areas pertaining to outcome prediction, anticipation and motor planning (Alexander & Brown, 2011; Thickbroom et al., 2000; Wolff et al., 2020) either showed no response (e.g., no activation in dorsal thalamus, supplementary motor areas) or demonstrated deactivation instead of activation (e.g., mPFC deactivation) during meditation relative to rest.

## 5 Conclusion

This is one of the first studies investigating meditation using ultra-high strength 7 Tesla fMRI. We found that during focused attention meditation relative to non-focused rest, a small sample of beginner meditators reliably attenuated key Default-mode network areas (i.e., antero-medial prefrontal cortex, posterior cingulate cortex, and precuneus) that typically subserve self-referential processing and mind-wandering. Additionally, we also observed significant deactivations in specific thalamic nuclei and visual areas suggesting perceptual decoupling during focused attention meditation relative to rest. Notably, these findings were significant after controlling for various physiological response fluctuations, head motion, multiple comparisons, overall baseline dispositional mindfulness, alertness during meditation and effort during meditation. Physiological relaxation, indexed by significant reduction in heart and breathing rates along with significant increase in breathing depth, was observed during meditation relative to rest. This lends additional support that individuals likely achieved meditative states as expected during the 7 Tesla fMRI acquisition. Significant brain findings were altered when correction for physiological responses was excluded. We also observed significant longitudinal changes in self-reported state mindfulness following a single session of fMRI meditation. However, these behavioural findings are potentially confounded by the small sample and possible placebo effects. Overall, this pilot study establishes the feasibility and utility of investigating focused attention meditation with beginner meditators using ultra-high strength 7 Tesla fMRI. We recommend the inclusion of physiological control and behavioural assessments in future larger neuroimaging investigations of meditation.

## Acknowledgements

The authors acknowledge the facilities (7 Tesla MRI scanner), scientific expertise and technical assistance (Rebecca Glarin and Braden Thai) of the National Imaging Facility, a National Collaborative Research Infrastructure Strategy (NCRIS) capability, at the Melbourne Brain Centre Imaging Unit, The University of Melbourne.

## Notes

Funding This research did not receive any specific grant from funding agencies in the public, commercial, or not-for-profit sectors. SG is supported by Australian Research Training Program (RTP) scholarship. AZ is supported by NHMRC Senior Research fellowship (APP1118153) and the Rebecca L. Cooper Fellowship. VL is supported by an AI and Val Rosenstrauss Senior Research Fellowship. NTVD is supported by the Contemplative Studies Centre, founded by a philanthropic gift from the Three Springs Foundation Pty Ltd.

### Competing Interest Statement

The authors have declared no competing interest.

## References

Ahani, A., Wahbeh, H., Miller, M., Nezamfar, H., Erdogmus, D., & Oken, B. (2013, 6-8 Nov. 2013). Change in physiological signals during mindfulness meditation. Paper presented at the 2013 6th International IEEE/EMBS Conference on Neural Engineering (NER).

Alberton, B. A. V., Nichols, T. E., Gamba, H. R., & Winkler, A. M. (2020). Multiple testing correction over contrasts for brain imaging. Neuroimage, 216, 116760. doi:https://doi.org/10.1016/j.neuroimage.2020.116760

Alexander, W. H., & Brown, J. W. (2011). Medial prefrontal cortex as an action-outcome predictor. Nature Neuroscience, 14(10), 1338–1344. doi:10.1038/nn.2921

Andersson, J. L. R., Skare, S., & Ashburner, J. (2003). How to correct susceptibility distortions in spin-echo echo-planar images: application to diffusion tensor imaging. Neuroimage, 20(2), 870–888. doi:https://doi.org/10.1016/S1053-8119(03)00336-7

Andrews-Hanna, J. R., Reidler, J. S., Sepulcre, J., Poulin, R., & Buckner, R. L. (2010). Functional-Anatomic Fractionation of the Brain’s Default Network. Neuron, 65(4), 550–562. doi:https://doi.org/10.1016/j.neuron.2010.02.005

Arch, J. J., & Craske, M. G. (2006). Mechanisms of mindfulness: Emotion regulation following a focused breathing induction. Behaviour Research and Therapy, 44(12), 1849–1858. doi:https://doi.org/10.1016/j.brat.2005.12.007

Avants, B. B., Tustison, N., & Song, G. (2009). Advanced normalization tools (ANTS). Insight j, 2(365), 1–35.

Baer, R. A., Smith, G. T., Hopkins, J., Krietemeyer, J., & Toney, L. (2006). Using self-report assessment methods to explore facets of mindfulness. Assessment, 13(1), 27–45. doi:10.1177/1073191105283504

Baron Short, E., Kose, S., Mu, Q., Borckardt, J., Newberg, A., George, M. S., & Kozel, F. A. (2010). Regional brain activation during meditation shows time and practice effects: an exploratory FMRI study. Evidence-based complementary and alternative medicine : eCAM, 7(1), 121–127. doi:10.1093/ecam/nem163

Beisteiner, R., Robinson, S., Wurnig, M., Hilbert, M., Merksa, K., Rath, J., … Geißler, A. (2011). Clinical fMRI: Evidence for a 7T benefit over 3T. Neuroimage, 57(3), 1015–1021. doi:https://doi.org/10.1016/j.neuroimage.2011.05.010

Birn, R. M., Diamond, J. B., Smith, M. A., & Bandettini, P. A. (2006). Separating respiratory-variation-related fluctuations from neuronal-activity-related fluctuations in fMRI. Neuroimage, 31(4), 1536–1548. doi:10.1016/j.neuroimage.2006.02.048

Birn, R. M., Murphy, K., Handwerker, D. A., & Bandettini, P. A. (2009). fMRI in the presence of task-correlated breathing variations. Neuroimage, 47(3), 1092–1104. doi:10.1016/j.neuroimage.2009.05.030

Bishop, S. R., Lau, M., Shapiro, S., Carlson, L., Anderson, N. D., Carmody, J., … Velting, D. (2004). Mindfulness: a proposed operational definition. Clinical Psychology: Science and Practice, 11(3), 230.

Brandmeyer, T., & Delorme, A. (2021). Meditation and the Wandering Mind: A Theoretical Framework of Underlying Neurocognitive Mechanisms. Perspect Psychol Sci, 16(1), 39–66. doi:10.1177/1745691620917340

Britton, W. B., Lindahl, J. R., Cahn, B. R., Davis, J. H., & Goldman, R. E. (2014). Awakening is not a metaphor: the effects of Buddhist meditation practices on basic wakefulness. Annals of the New York Academy of Sciences, 1307(1), 64–81. doi:https://doi.org/10.1111/nyas.12279

Broderick, P. C. (2005). Mindfulness and Coping with Dysphoric Mood: Contrasts with Rumination and Distraction. Cognitive Therapy and Research, 29(5), 501–510. doi:10.1007/s10608-005-3888-0

Cavanna, A. E., & Trimble, M. R. (2006). The precuneus: a review of its functional anatomy and behavioural correlates. Brain, 129(3), 564–583.

Chang, C., Cunningham, J. P., & Glover, G. H. (2009). Influence of heart rate on the BOLD signal: the cardiac response function. Neuroimage, 44(3), 857–869. doi:10.1016/j.neuroimage.2008.09.029

Creswell, J. D. (2017). Mindfulness Interventions. Annual Review of Psychology, 68(1), 491–516. doi:10.1146/annurev-psych-042716-051139

Delmonte, M. M. (1984). Physiological responses during meditation and rest. Biofeedback and Self-regulation, 9(2), 181–200. doi:10.1007/BF00998833

Dickenson, J., Berkman, E. T., Arch, J., & Lieberman, M. D. (2013). Neural correlates of focused attention during a brief mindfulness induction. Soc Cogn Affect Neurosci, 8(1), 40–47. doi:10.1093/scan/nss030

Ditto, B., Eclache, M., & Goldman, N. (2006). Short-term autonomic and cardiovascular effects of mindfulness body scan meditation. Ann Behav Med, 32(3), 227–234. doi:10.1207/s15324796abm3203_9

Doll, A., Hölzel, B. K., Mulej Bratec, S., Boucard, C. C., Xie, X., Wohlschläger, A. M., & Sorg, C. (2016). Mindful attention to breath regulates emotions via increased amygdala-prefrontal cortex connectivity. Neuroimage, 134, 305–313. doi:10.1016/j.neuroimage.2016.03.041

Engström, M., Willander, J., & Simon, R. (2022). A Review of the Methodology, Taxonomy, and Definitions in Recent fMRI Research on Meditation. Mindfulness, 13(3), 541–555. doi:10.1007/s12671-021-01782-7

Farb, N. A., Segal, Z. V., & Anderson, A. K. (2013). Mindfulness meditation training alters cortical representations of interoceptive attention. Soc Cogn Affect Neurosci, 8(1), 15–26. doi:10.1093/scan/nss066

Farias, M., Maraldi, E., Wallenkampf, K., & Lucchetti, G. (2020). Adverse events in meditation practices and meditation-based therapies: a systematic review. Acta Psychiatrica Scandinavica, 142(5), 374–393.

Feruglio, S., Matiz, A., Pagnoni, G., Fabbro, F., & Crescentini, C. (2021). The Impact of Mindfulness Meditation on the Wandering Mind: a Systematic Review. Neurosci Biobehav Rev, 131, 313–330. doi:10.1016/j.neubiorev.2021.09.032

Fox, K. C., Dixon, M. L., Nijeboer, S., Girn, M., Floman, J. L., Lifshitz, M., Christoff, K. (2016). Functional neuroanatomy of meditation: A review and meta-analysis of 78 functional neuroimaging investigations. Neurosci Biobehav Rev, 65, 208–228. doi:10.1016/j.neubiorev.2016.03.021

Fox, K. C., Spreng, R. N., Ellamil, M., Andrews-Hanna, J. R., & Christoff, K. (2015). The wandering brain: meta-analysis of functional neuroimaging studies of mind-wandering and related spontaneous thought processes. Neuroimage, 111, 611–621. doi:10.1016/j.neuroimage.2015.02.039

Frewen, P. A., Evans, E. M., Maraj, N., Dozois, D. J. A., & Partridge, K. (2008). Letting Go: Mindfulness and Negative Automatic Thinking. Cognitive Therapy and Research, 32(6), 758–774. doi:10.1007/s10608-007-9142-1

Galante, J., Friedrich, C., Dawson, A. F., Modrego-Alarcón, M., Gebbing, P., Delgado-Suárez, I., … Jones, P. B. (2021). Mindfulness-based programmes for mental health promotion in adults in nonclinical settings: A systematic review and meta-analysis of randomised controlled trials. PLoS Med, 18(1), e1003481. doi:10.1371/journal.pmed.1003481

Ganesan, S., Beyer, E., Moffat, B., Van Dam, N. T., Lorenzetti, V., & Zalesky, A. (2022a). Focused attention meditation in healthy adults: A systematic review and meta-analysis of cross-sectional functional MRI studies. Neuroscience & Biobehavioral Reviews, 141, 104846. doi:https://doi.org/10.1016/j.neubiorev.2022.104846

Ganesan, S., Lv, J., & Zalesky, A. (2022b). Multi-timepoint pattern analysis: Influence of personality and behavior on decoding context-dependent brain connectivity dynamics. Human Brain Mapping, 43(4), 1403–1418. doi:https://doi.org/10.1002/hbm.25732

Gizewski, E. R., de Greiff, A., Maderwald, S., Timmann, D., Forsting, M., & Ladd, M. E. (2007). fMRI at 7 T: Whole-brain coverage and signal advantages even infratentorially? Neuroimage, 37(3), 761–768. doi:https://doi.org/10.1016/j.neuroimage.2007.06.005

Glasser, M. F., Coalson, T. S., Robinson, E. C., Hacker, C. D., Harwell, J., Yacoub, E., Van Essen, D. C. (2016). A multi-modal parcellation of human cerebral cortex. Nature, 536(7615), 171–178. doi:10.1038/nature18933

Glasser, M. F., Sotiropoulos, S. N., Wilson, J. A., Coalson, T. S., Fischl, B., Andersson, J. L., … Jenkinson, M. (2013). The minimal preprocessing pipelines for the Human Connectome Project. Neuroimage, 80, 105–124. doi:10.1016/j.neuroimage.2013.04.127

Glover, G. H., Li, T. Q., & Ress, D. (2000). Image-based method for retrospective correction of physiological motion effects in fMRI: RETROICOR. Magn Reson Med, 44(1), 162–167. doi:10.1002/1522-2594(200007)44:1<162::aid-mrm23>3.0.co;2-e

Grill-Spector, K., Kourtzi, Z., & Kanwisher, N. (2001). The lateral occipital complex and its role in object recognition. Vision research, 41(10-11), 1409–1422.

Hale, J. R., Brookes, M. J., Hall, E. L., Zumer, J. M., Stevenson, C. M., Francis, S. T., & Morris, P. G. (2010). Comparison of functional connectivity in default mode and sensorimotor networks at 3 and 7T. Magnetic Resonance Materials in Physics, Biology and Medicine, 23(5), 339–349. doi:10.1007/s10334-010-0220-0

Harrison, S. J., Bianchi, S., Heinzle, J., Stephan, K. E., Iglesias, S., & Kasper, L. (2021). A Hilbert-based method for processing respiratory timeseries. Neuroimage, 230, 117787. doi:10.1016/j.neuroimage.2021.117787

Hasenkamp, W., Wilson-Mendenhall, C. D., Duncan, E., & Barsalou, L. W. (2012). Mind wandering and attention during focused meditation: a fine-grained temporal analysis of fluctuating cognitive states. Neuroimage, 59(1), 750–760. doi:10.1016/j.neuroimage.2011.07.008

Hölzel, B. K., Ott, U., Hempel, H., Hackl, A., Wolf, K., Stark, R., & Vaitl, D. (2007). Differential engagement of anterior cingulate and adjacent medial frontal cortex in adept meditators and non-meditators. Neurosci Lett, 421(1), 16–21. doi:10.1016/j.neulet.2007.04.074

Hwang, K., Bertolero, M. A., Liu, W. B., & Esposito, M. (2017). The Human Thalamus Is an Integrative Hub for Functional Brain Networks. The Journal of Neuroscience, 37(23), 5594. doi:10.1523/JNEUROSCI.0067-17.2017

Jenkinson, M., Bannister, P., Brady, M., & Smith, S. (2002). Improved optimization for the robust and accurate linear registration and motion correction of brain images. Neuroimage, 17(2), 825–841. doi:10.1016/s1053-8119(02)91132-8

Jenkinson, M., & Smith, S. (2001). A global optimisation method for robust affine registration of brain images. Med Image Anal, 5(2), 143–156. doi:10.1016/s1361-8415(01)00036-6

Jha, A. P., Krompinger, J., & Baime, M. J. (2007). Mindfulness training modifies subsystems of attention. Cognitive, Affective, & Behavioral Neuroscience, 7(2), 109–119.

Johnson, S., Gur, R. M., David, Z., & Currier, E. (2015). One-Session Mindfulness Meditation: A Randomized Controlled Study of Effects on Cognition and Mood. Mindfulness, 6(1), 88–98. doi:10.1007/s12671-013-0234-6

k, F. W., s, A. A. M., Chee, M. W. L., & Lim, J. (2018). Towards an Objective Measure of Mindfulness: Replicating and Extending the Features of the Breath-Counting Task. Mindfulness (N Y), 9(5), 1402–1410. doi:10.1007/s12671-017-0880-1

Kasper, L., Bollmann, S., Diaconescu, A. O., Hutton, C., Heinzle, J., Iglesias, S., … Stephan, K. E. (2017). The PhysIO Toolbox for Modeling Physiological Noise in fMRI Data. Journal of Neuroscience Methods, 276, 56–72. doi:https://doi.org/10.1016/j.jneumeth.2016.10.019

Kim, H. C., Tegethoff, M., Meinlschmidt, G., Stalujanis, E., Belardi, A., Jo, S., Lee, J. H. (2019). Mediation analysis of triple networks revealed functional feature of mindfulness from real-time fMRI neurofeedback. Neuroimage, 195, 409–432. doi:10.1016/j.neuroimage.2019.03.066

Laukkonen, R. E., & Slagter, H. A. (2021). From many to (n)one: Meditation and the plasticity of the predictive mind. Neuroscience & Biobehavioral Reviews, 128, 199–217. doi:https://doi.org/10.1016/j.neubiorev.2021.06.021

Leech, R., & Sharp, D. J. (2014). The role of the posterior cingulate cortex in cognition and disease. Brain, 137(Pt 1), 12–32. doi:10.1093/brain/awt162

Leech, R., & Smallwood, J. (2019). The posterior cingulate cortex: Insights from structure and function. Handbook of clinical neurology, 166, 73–85.

Levinson, D. B., Stoll, E. L., Kindy, S. D., Merry, H. L., & Davidson, R. J. (2014). A mind you can count on: validating breath counting as a behavioral measure of mindfulness. Frontiers in psychology, 5, 1202–1202. doi:10.3389/fpsyg.2014.01202

Li, X., Morgan, P. S., Ashburner, J., Smith, J., & Rorden, C. (2016). The first step for neuroimaging data analysis: DICOM to NIfTI conversion. J Neurosci Methods, 264, 47–56. doi:10.1016/j.jneumeth.2016.03.001

Lieberman, M. D., Straccia, M. A., Meyer, M. L., Du, M., & Tan, K. M. (2019). Social, self, (situational), and affective processes in medial prefrontal cortex (MPFC): Causal, multivariate, and reverse inference evidence. Neurosci Biobehav Rev, 99, 311–328. doi:10.1016/j.neubiorev.2018.12.021

Lomas, T., Cartwright, T., Edginton, T., & Ridge, D. (2015). A qualitative analysis of experiential challenges associated with meditation practice. Mindfulness, 6(4), 848–860.

Lutz, A., Slagter, H. A., Dunne, J. D., & Davidson, R. J. (2008). Attention regulation and monitoring in meditation. Trends in cognitive sciences, 12(4), 163–169. doi:10.1016/j.tics.2008.01.005

Manna, A., Raffone, A., Perrucci, M. G., Nardo, D., Ferretti, A., Tartaro, A., Romani, G. L. (2010). Neural correlates of focused attention and cognitive monitoring in meditation. Brain Res Bull, 82(1-2), 46–56. doi:10.1016/j.brainresbull.2010.03.001

Matko, K., Ott, U., & Sedlmeier, P. (2021). What Do Meditators Do When They Meditate? Proposing a Novel Basis for Future Meditation Research. Mindfulness, 12(7), 1791–1811. doi:10.1007/s12671-021-01641-5

Matko, K., & Sedlmeier, P. (2019). What Is Meditation? Proposing an Empirically Derived Classification System. Frontiers in psychology, 10(2276). doi:10.3389/fpsyg.2019.02276

May, L. M., Reinka, M. A., Tipsord, J. M., Felver, J. C., & Berkman, E. T. (2016). Parenting an Early Adolescent: a Pilot Study Examining Neural and Relationship Quality Changes of a Mindfulness Intervention. Mindfulness, 7(5), 1203–1213. doi:10.1007/s12671-016-0563-3

Melis, M., Schroyen, G., Pollefeyt, J., Raes, F., Smeets, A., Sunaert, S., … Van der Gucht, K. (2022). The Impact of Mindfulness-Based Interventions on Brain Functional Connectivity: a Systematic Review. Mindfulness, 13(8), 1857–1875. doi:10.1007/s12671-022-01919-2

Moeller, S., Yacoub, E., Olman, C. A., Auerbach, E., Strupp, J., Harel, N., & Uğurbil, K. (2010). Multiband multislice GE-EPI at 7 tesla, with 16-fold acceleration using partial parallel imaging with application to high spatial and temporal whole-brain fMRI. Magn Reson Med, 63(5), 1144–1153. doi:10.1002/mrm.22361

Mohan, A., Sharma, R., & Bijlani, R. L. (2011). Effect of Meditation on Stress-Induced Changes in Cognitive Functions. The Journal of Alternative and Complementary Medicine, 17(3), 207–212. doi:10.1089/acm.2010.0142

Mooneyham, B. W., Mrazek, M. D., Mrazek, A. J., Mrazek, K. L., Phillips, D. T., & Schooler, J. W. (2017). States of Mind: Characterizing the Neural Bases of Focus and Mind-wandering through Dynamic Functional Connectivity. J Cogn Neurosci, 29(3), 495–506. doi:10.1162/jocn_a_01066

Mrazek, M. D., Smallwood, J., & Schooler, J. W. (2012). Mindfulness and mind-wandering: finding convergence through opposing constructs. Emotion, 12(3), 442.

Müller, C., Dubiel, D., Kremeti, E., Lieb, M., Streicher, E., Siakir Oglou, N., … Karbach, J. (2021). Effects of a Single Physical or Mindfulness Intervention on Mood, Attention, and Executive Functions: Results from two Randomized Controlled Studies in University Classes. Mindfulness, 12(5), 1282–1293. doi:10.1007/s12671-021-01601-z

Nichols, T. E., & Holmes, A. P. (2002). Nonparametric permutation tests for functional neuroimaging: a primer with examples. Hum Brain Mapp, 15(1), 1–25. doi:10.1002/hbm.1058

O’Brien, K. R., Kober, T., Hagmann, P., Maeder, P., Marques, J., Lazeyras, F., … Roche, A. (2014). Robust T1-weighted structural brain imaging and morphometry at 7T using MP2RAGE. PLoS One, 9(6), e99676. doi:10.1371/journal.pone.0099676

Pohmann, R., Speck, O., & Scheffler, K. (2016). Signal-to-noise ratio and MR tissue parameters in human brain imaging at 3, 7, and 9.4 tesla using current receive coil arrays. Magn Reson Med, 75(2), 801–809. doi:10.1002/mrm.25677

Prescott, S. A., & Ratté, S. (2017). Chapter 23 - Somatosensation and Pain. In P. M. Conn (Ed.), Conn’s Translational Neuroscience (pp. 517–539). San Diego: Academic Press.

Raichle, M. E., MacLeod, A. M., Snyder, A. Z., Powers, W. J., Gusnard, D. A., & Shulman, G. L. (2001). A default mode of brain function. Proc Natl Acad Sci U S A, 98(2), 676–682. doi:10.1073/pnas.98.2.676

Ritskes, R., Ritskes-Hoitinga, A., Stødkilde-Jørgensen, A., Bærentsen, K. B., & Hartmann, T. (2004). MRI scanning during Zen meditation: the picture of enlightenment. The relevance of the wisdom traditions in contemporary society: the challenge to psychology, 195–198.

Scheibner, H. J., Bogler, C., Gleich, T., Haynes, J. D., & Bermpohl, F. (2017). Internal and external attention and the default mode network. Neuroimage, 148, 381–389. doi:10.1016/j.neuroimage.2017.01.044

Schlosser, M., Sparby, T., Vörös, S., Jones, R., & Marchant, N. L. (2019). Unpleasant meditation-related experiences in regular meditators: Prevalence, predictors, and conceptual considerations. PLoS One, 14(5), e0216643. doi:10.1371/journal.pone.0216643

Seghier, M. L. (2013). The angular gyrus: multiple functions and multiple subdivisions. Neuroscientist, 19(1), 43–61. doi:10.1177/1073858412440596

Sezer, I., Pizzagalli, D. A., & Sacchet, M. D. (2022). Resting-state fMRI functional connectivity and mindfulness in clinical and non-clinical contexts: A review and synthesis. Neuroscience & Biobehavioral Reviews, 135, 104583. doi:https://doi.org/10.1016/j.neubiorev.2022.104583

Shapiro, S. L., & Walsh, R. (2003). An analysis of recent meditation research and suggestions for future directions. The Humanistic Psychologist, 31(2-3), 86–114.

Soni, R., & Muniyandi, M. (2019). Breath Rate Variability: A Novel Measure to Study the Meditation Effects. Int J Yoga, 12(1), 45–54. doi:10.4103/ijoy.IJOY_27_17

Spielberger, C., Gorsuch, R., Lushene, R., Vagg, P. R., & Jacobs, G. (1983). Manual for the State-Trait Anxiety Inventory (Form Y1 – Y2) (Vol. IV).

Swenson, R. S., & Gulledge, A. T. (2017). Chapter 12 - The Cerebral Cortex. In P. M. Conn (Ed.), Conn’s Translational Neuroscience (pp. 263–288). San Diego: Academic Press.

Tanay, G., & Bernstein, A. (2013). State Mindfulness Scale (SMS): development and initial validation. Psychol Assess, 25(4), 1286–1299. doi:10.1037/a0034044

Theysohn, N., Qin, S., Maderwald, S., Poser, B. A., Theysohn, J. M., Ladd, M. E., Tendolkar, I. (2013). Memory-Related Hippocampal Activity Can Be Measured Robustly Using fMRI at 7 Tesla. Journal of Neuroimaging, 23(4), 445–451. doi:https://doi.org/10.1111/jon.12036

Thickbroom, G. W., Byrnes, M. L., Sacco, P., Ghosh, S., Morris, I. T., & Mastaglia, F. L. (2000). The role of the supplementary motor area in externally timed movement: the influence of predictability of movement timing. Brain Research, 874(2), 233–241. doi:https://doi.org/10.1016/S0006-8993(00)02588-9

Torrisi, S., Chen, G., Glen, D., Bandettini, P. A., Baker, C. I., Reynolds, R., Ernst, M. (2018). Statistical power comparisons at 3T and 7T with a GO / NOGO task. Neuroimage, 175, 100–110. doi:https://doi.org/10.1016/j.neuroimage.2018.03.071

Trattnig, S., Springer, E., Bogner, W., Hangel, G., Strasser, B., Dymerska, B., Robinson, S. D. (2018). Key clinical benefits of neuroimaging at 7T. Neuroimage, 168, 477–489. doi:10.1016/j.neuroimage.2016.11.031

Trungpa, C. (2002). The myth of freedom and the way of meditation: Shambhala Publications.

Utevsky, A. V., Smith, D. V., & Huettel, S. A. (2014). Precuneus Is a Functional Core of the Default-Mode Network. The Journal of Neuroscience, 34(3), 932. doi:10.1523/JNEUROSCI.4227-13.2014

Valentine, E. R., & Sweet, P. L. (1999). Meditation and attention: A comparison of the effects of concentrative and mindfulness meditation on sustained attention. Mental health, religion & culture, 2(1), 59–70.

Van Dam, N. T., Van Vugt, M. K., Vago, D. R., Schmalzl, L., Saron, C. D., Olendzki, A., … Gorchov, J. (2018). Mind the hype: A critical evaluation and prescriptive agenda for research on mindfulness and meditation. Perspectives on Psychological Science, 13(1), 36–61.

Viessmann, O., & Polimeni, J. R. (2021). High-resolution fMRI at 7 Tesla: challenges, promises and recent developments for individual-focused fMRI studies. Curr Opin Behav Sci, 40, 96–104. doi:10.1016/j.cobeha.2021.01.011

Weiner, K. S., & Zilles, K. (2016). The anatomical and functional specialization of the fusiform gyrus. Neuropsychologia, 83, 48–62. doi:https://doi.org/10.1016/j.neuropsychologia.2015.06.033

Weng, H. Y., Lewis-Peacock, J. A., Hecht, F. M., Uncapher, M. R., Ziegler, D. A., Farb, N. A. S., … Gazzaley, A. (2020). Focus on the Breath: Brain Decoding Reveals Internal States of Attention During Meditation. Front Hum Neurosci, 14, 336. doi:10.3389/fnhum.2020.00336

Wenk-Sormaz, H. (2005). Meditation can reduce habitual responding. Altern Ther Health Med, 11(2), 42–58.

Winkler, A. M., Ridgway, G. R., Douaud, G., Nichols, T. E., & Smith, S. M. (2016). Faster permutation inference in brain imaging. Neuroimage, 141, 502–516. doi:https://doi.org/10.1016/j.neuroimage.2016.05.068

Winkler, A. M., Ridgway, G. R., Webster, M. A., Smith, S. M., & Nichols, T. E. (2014). Permutation inference for the general linear model. Neuroimage, 92, 381–397. doi:https://doi.org/10.1016/j.neuroimage.2014.01.060

Wolff, S., Kohrs, C., Angenstein, N., & Brechmann, A. (2020). Dorsal posterior cingulate cortex encodes the informational value of feedback in human–computer interaction. Scientific reports, 10(1), 13030. doi:10.1038/s41598-020-68300-y

Woolrich, M. W., Ripley, B. D., Brady, M., & Smith, S. M. (2001). Temporal Autocorrelation in Univariate Linear Modeling of FMRI Data. Neuroimage, 14(6), 1370–1386. doi:https://doi.org/10.1006/nimg.2001.0931

Yamaya, N., Tsuchiya, K., Takizawa, I., Shimoda, K., Kitazawa, K., & Tozato, F. (2021). Effect of one-session focused attention meditation on the working memory capacity of meditation novices: A functional near-infrared spectroscopy study. Brain and behavior, 11(8), e2288. doi:https://doi.org/10.1002/brb3.2288

Young, K. S., van der Velden, A. M., Craske, M. G., Pallesen, K. J., Fjorback, L., Roepstorff, A., & Parsons, C. E. (2018). The impact of mindfulness-based interventions on brain activity: A systematic review of functional magnetic resonance imaging studies. Neuroscience and Biobehavioral Reviews, 84, 424–433. doi:10.1016/j.neubiorev.2017.08.003

