## Supplementary material for "Meditation attenuates Default-mode activity: a pilot study using ultra-high strength MRI"

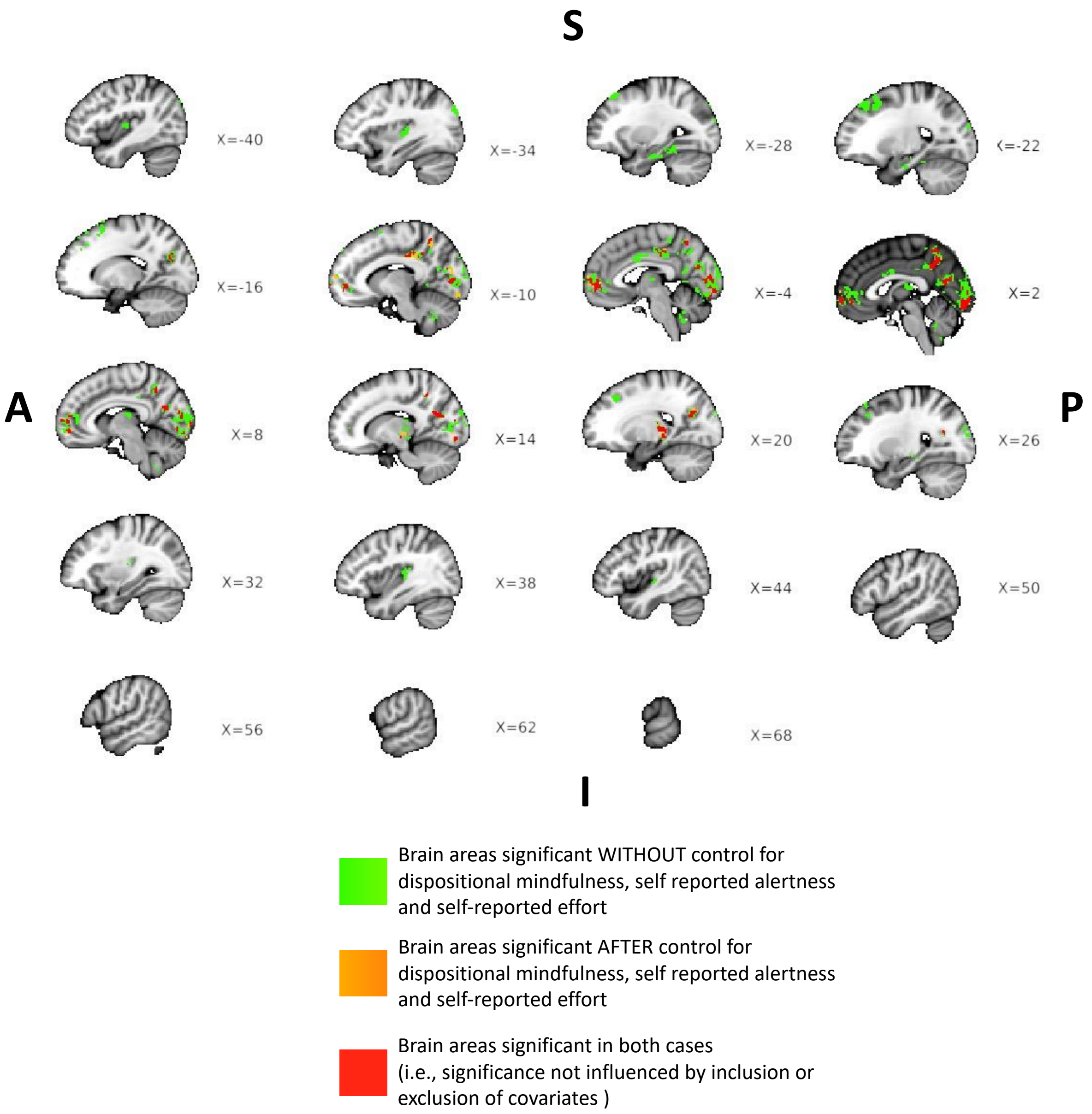


***Fig. S1*** *Section of volumetric sagittal brain slices along with corresponding x-coordinates displaying the differences in the extent of significantly de-activated brain areas during meditation relative to rest, when group-level covariates (i.e., dispositional mindfulness, self-reported alertness during meditation, self-reported effort during meditation) were included vs. excluded. Brain areas coloured green were only significant when covariates were excluded. Brain areas coloured orange were only significant after inclusion of the covariates. Brain*

*areas whose significance was not influenced by inclusion or exclusion of covariates are shown in red. All significant brain areas survived FWE correction for multiple comparisons across clusters (as reported in main manuscript). S – superior/dorsal, A – anterior, P – posterior, I – inferior/ventral.*

When not controlling for inter-individual variability in overall baseline dispositional mindfulness (overall FFMQ scores), self-reported alertness during meditation and self-reported effort during meditation, we found that the extent of significant deactivation clusters for meditation relative to rest was slightly more diffuse (green and red areas in Fig. S1). Particularly, the extent of deactivation was more diffuse in the visual areas, thalamus, posterior cingulate cortex (PCC), precuneus and medial prefrontal cortex (mPFC) (which also included dorsomedial PFC). Additional deactivation was also found in the anterior cingulate cortex (ACC), posterior insula, hippocampal areas, and cerebellum. These regions have been implicated previously in mind-wandering distraction and spontaneous thought. Specifically, parts of posterior insula may be involved in the somatic experience of mind-wandering, hinted by its reported activation during mind-wandering processes (Fox et al., 2015) and reported deactivation during focused attention meditation (Hasenkamp et al., 2012; Manna et al., 2010). Deactivation of hippocampal areas during meditation relative to rest likely result from attenuation of memory recall associated with default-mode processing (Buckner & Carroll, 2007; Fox et al., 2015; Hasenkamp et al., 2012). Deactivation within parts of ACC during meditation relative to rest may be explained by its relevance to goal-directed cognition (e.g., thoughts about meditating) (Fox et al., 2015; Hasenkamp et al., 2012), as well as mind-wandering without meta-awareness (Christoff et al., 2009). Finally, although some studies report the involvement of cerebellum in focused attention meditation (Hasenkamp et al., 2012; Miyoshi et al., 2019), the role of cerebellum in meditation is yet to be explored. The observed deactivation of cerebellum during meditation relative to rest may potentially underpin attenuation of abstract mentalizing and social cognition (Van Overwalle et al., 2014), alongside deactivation of Default-mode processing.

On the other hand, controlling for these inter-individual variables increased the specificity of deactivations, confining them to specific sub areas within the visual cortex, PCC, mPFC and thalamus (orange and red areas in Fig. S1; see main text for more details). Additionally, the deactivations within posterior insula, ACC, hippocampal areas and cerebellum lost statistical significance. In other words, the significance of deactivations in these regions was likely influenced by level of variations in overall dispositional mindfulness, subjective effort during meditation and subjective arousal during meditation. Previous studies have found associations between meditation experience and fMRI activity in posterior insula (Farb et al., 2013; Hasenkamp et al., 2012; Manna et al., 2010), para-hippocampus, ACC and cerebellar regions (Hasenkamp et al., 2012). Similarly, other brain areas within networks such as Default-mode and Executive Control can be influenced by inter-individual variability in dispositional mindfulness levels (see Ganesan et al. (2022) for detailed review), as well as level of arousal and effort during meditation task among beginners (see Britton et al. (2014) for a detailed review). Moreover, meditators with greater mindfulness ability potentially demonstrate greater neural efficiency during meditation (Brefczynski-Lewis et al., 2007; Escrichs et al., 2019; Hiroyasu & Hiwa, 2017; Manna et al., 2010). Taken together, it is possible that inter-individual variations in dispositional mindfulness, arousal and effort may have influenced the extent and areas of deactivations observed and consequently, controlling for these factors increased the precision of deactivations in our study. However, there is a definitive need for these findings to be validated in bigger samples.

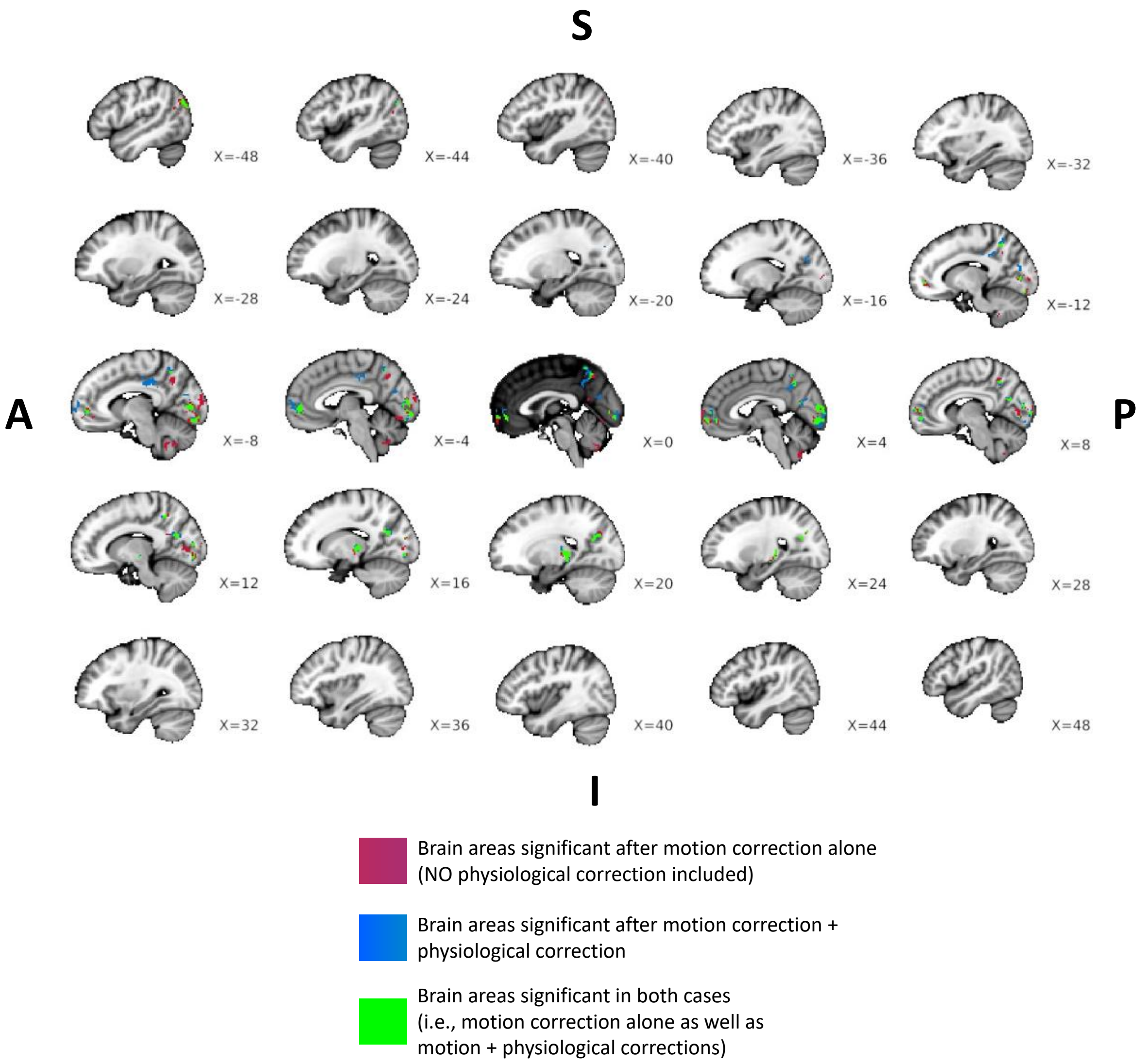


***Fig. S2*** *Section of volumetric sagittal brain slices along with corresponding x-coordinates displaying the differences in the extent of significantly de-activated brain areas during meditation relative to rest, when physiological artifact correction was included vs. excluded. Brain areas coloured maroon were only significant when physiological correction was excluded. Brain areas coloured blue were only significant after inclusion of physiological correction. Brain areas whose significance was not affected by physiological correction are shown in green. All significant brain areas survived FWE correction for multiple comparisons across clusters (as reported in main manuscript). Note that all significant brain areas shown*

*here were obtained after controlling for overall dispositional mindfulness, and self-report effort and self-report alertness during meditation. S – superior/dorsal, A – anterior, P – posterior, I – inferior/ventral.*

As shown by the maroon and blue areas in Fig. S2, the inclusion vs. exclusion of physiological signal control in fMRI analysis impacts the extent of significance among deactivated mid-line cortical areas (including PCC, mPFC and precuneus) during meditation relative to rest. Physiological artifact correction can influence BOLD fMRI signal, especially along the mid-line cortical areas near large blood vessels, such as PCC and precuneus. In general, default-mode network areas are likely impacted by task-induced as well as respiration-induced blood flow changes due to the nearby large vasculatures (Birn et al., 2006). Furthermore, these physiological confound effects can be greater when fMRI task conditions closely associate with physiological response fluctuations (Birn et al., 2009), like with meditation (Ahani et al., 2013; Delmonte, 1984; Ditto et al., 2006; Soni & Muniyandi, 2019). Therefore, it is important that future neuroimaging studies investigating meditation report fMRI results that have been controlled for physiological confounds.

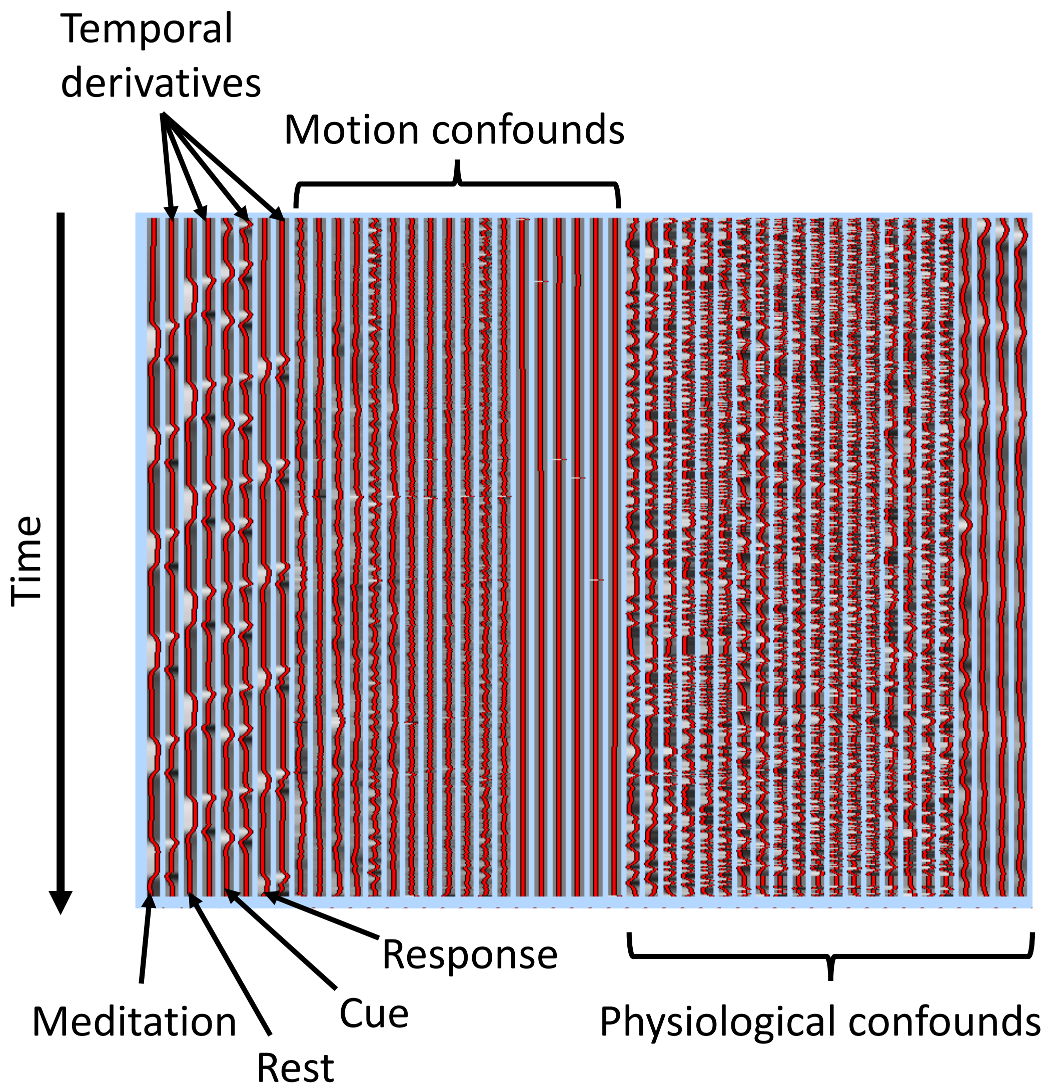


***Fig. S3*** *Visual depiction of the design matrix used in the first-level general linear modelling (GLM) analysis of every fMRI run. Time courses (shown in red) of predictors modelling the HRF-convolved main conditions of interest (i.e., meditation, rest, cue, response), their respective temporal derivatives, head motion artifacts, and physiological artifacts constitute the first-level design matrix. The y-axis represents time of fMRI run from start to finish.*

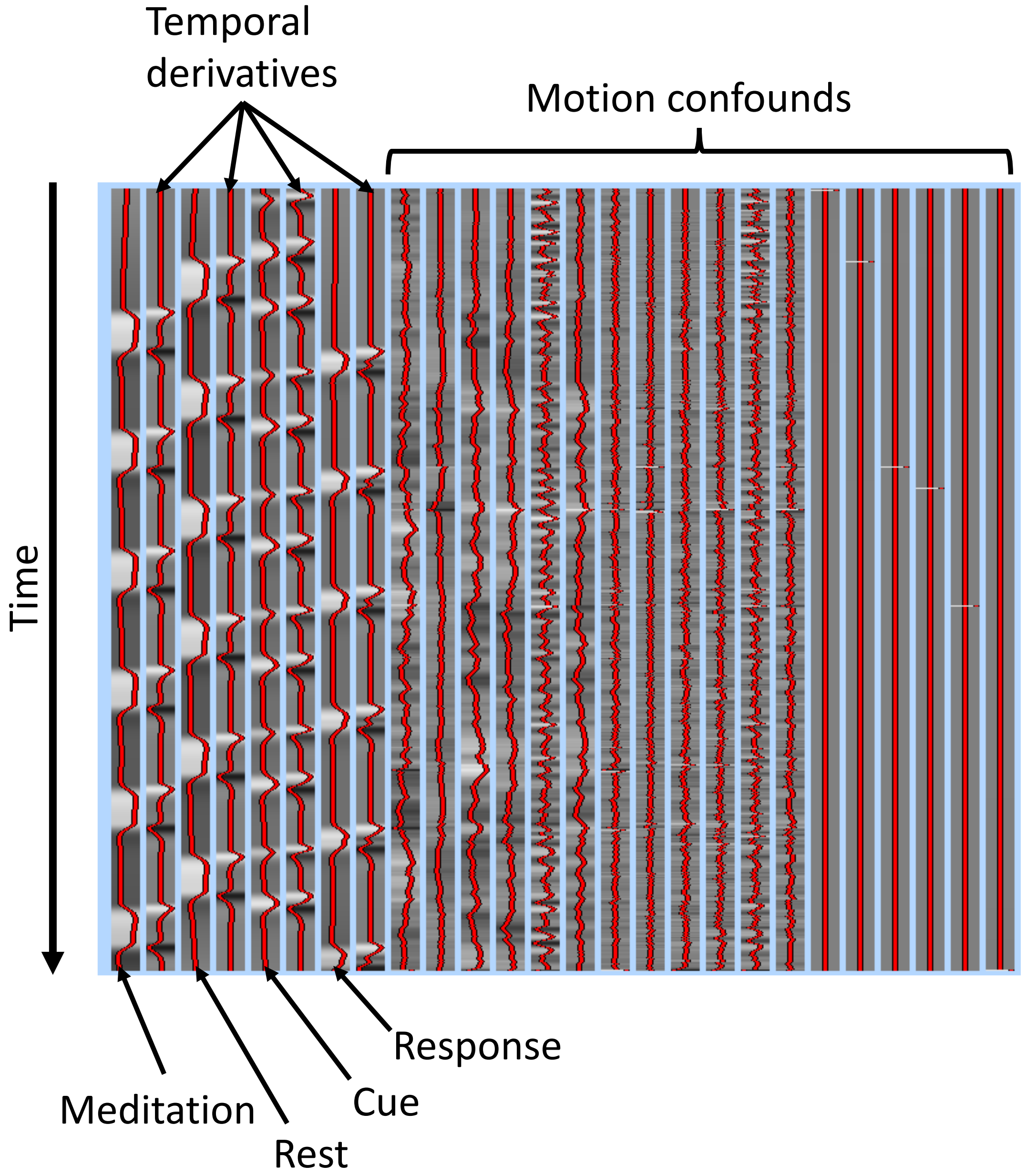


***Fig. S4** Visual depiction of the design matrix used in the first-level general linear modelling (GLM) analysis when physiological correction was excluded. Time courses (shown in red) of predictors modelling the HRF-convolved main conditions of interest (i.e., meditation, rest, cue, response), their respective temporal derivatives, and head motion artifacts constitute this first-level design matrix when physiological correction was entirely excluded. The y-axis represents time of fMRI run from start to finish.*

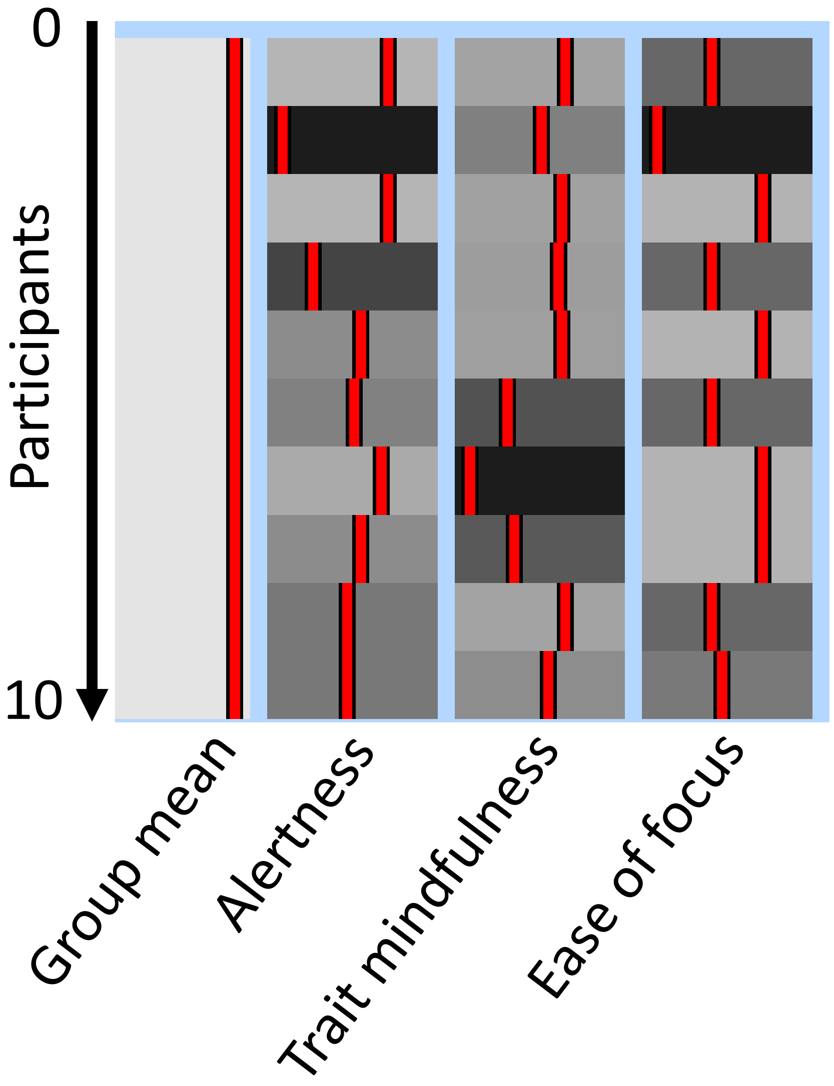


***Fig. S5*** *Visual depiction of the design matrix with covariates used in the third-level (group level) general linear modelling (GLM) analysis. The first column models the average response during meditation relative to rest across the 10 participants. The remaining columns model the inter-individual variability in self-reported in-scanner alertness, trait or dispositional mindfulness (overall FFMQ score), and self-reported in-scanner effort. The y-axis represents participants. Note that a group-level design matrix that excludes the covariates (as detailed in Fig. S1) would comprise only the 'group mean' column shown here.*

### References

Ahani, A., Wahbeh, H., Miller, M., Nezamfar, H., Erdogmus, D., & Oken, B. (2013, 6-8 Nov. 2013). *Change in physiological signals during mindfulness meditation.* Paper presented at the 2013 6th International IEEE/EMBS Conference on Neural Engineering (NER).

Birn, R. M., Diamond, J. B., Smith, M. A., & Bandettini, P. A. (2006). Separating respiratory-variation-related fluctuations from neuronal-activity-related fluctuations in fMRI. *Neuroimage, 31*(4), 1536-1548. doi:10.1016/j.neuroimage.2006.02.048

Birn, R. M., Murphy, K., Handwerker, D. A., & Bandettini, P. A. (2009). fMRI in the presence of task-correlated breathing variations. *Neuroimage, 47*(3), 1092-1104. doi:10.1016/j.neuroimage.2009.05.030

Brefczynski-Lewis, J. A., Lutz, A., Schaefer, H. S., Levinson, D. B., & Davidson, R. J. (2007). Neural correlates of attentional expertise in long-term meditation practitioners. *Proc Natl Acad Sci U S A, 104*(27), 11483-11488. doi:10.1073/pnas.0606552104

Britton, W. B., Lindahl, J. R., Cahn, B. R., Davis, J. H., & Goldman, R. E. (2014). Awakening is not a metaphor: the effects of Buddhist meditation practices on basic wakefulness. *Annals of the New York Academy of Sciences, 1307*(1), 64-81. doi:https://doi.org/10.1111/nyas.12279

Buckner, R. L., & Carroll, D. C. (2007). Self-projection and the brain. *Trends in cognitive sciences, 11*(2), 49-57. doi:https://doi.org/10.1016/j.tics.2006.11.004

Christoff, K., Gordon, A. M., Smallwood, J., Smith, R., & Schooler, J. W. (2009). Experience sampling during fMRI reveals default network and executive system contributions to mind wandering. *Proc Natl Acad Sci U S A, 106*(21), 8719-8724. doi:10.1073/pnas.0900234106

Delmonte, M. M. (1984). Physiological responses during meditation and rest. *Biofeedback and Self-regulation, 9*(2), 181-200. doi:10.1007/BF00998833

Ditto, B., Eclache, M., & Goldman, N. (2006). Short-term autonomic and cardiovascular effects of mindfulness body scan meditation. *Ann Behav Med, 32*(3), 227-234. doi:10.1207/s15324796abm3203_9

Escrichs, A., Sanjuán, A., Atasoy, S., López-González, A., Garrido, C., Càmara, E., & Deco, G. (2019). Characterizing the Dynamical Complexity Underlying Meditation. *Front Syst Neurosci, 13*, 27. doi:10.3389/fnsys.2019.00027

Farb, N. A., Segal, Z. V., & Anderson, A. K. (2013). Mindfulness meditation training alters cortical representations of interoceptive attention. *Soc Cogn Affect Neurosci, 8*(1), 15-26. doi:10.1093/scan/nss066

Fox, K. C., Spreng, R. N., Ellamil, M., Andrews-Hanna, J. R., & Christoff, K. (2015). The wandering brain: meta-analysis of functional neuroimaging studies of mind-wandering and related spontaneous thought processes. *Neuroimage, 111*, 611-621. doi:10.1016/j.neuroimage.2015.02.039

Ganesan, S., Beyer, E., Moffat, B., Van Dam, N. T., Lorenzetti, V., & Zalesky, A. (2022). Focused attention meditation in healthy adults: A systematic review and meta-analysis of cross-sectional functional MRI studies. *Neuroscience & Biobehavioral Reviews, 141*, 104846. doi:https://doi.org/10.1016/j.neubiorev.2022.104846

Hasenkamp, W., Wilson-Mendenhall, C. D., Duncan, E., & Barsalou, L. W. (2012). Mind wandering and attention during focused meditation: a fine-grained temporal analysis of fluctuating cognitive states. *Neuroimage, 59*(1), 750-760. doi:10.1016/j.neuroimage.2011.07.008

Hiroyasu, T., & Hiwa, S. (2017). *Brain functional state analysis of mindfulness using graph theory and functional connectivity.* Paper presented at the AAAI Spring Symposium - Technical Report.

Manna, A., Raffone, A., Perrucci, M. G., Nardo, D., Ferretti, A., Tartaro, A., . . . Romani, G. L. (2010). Neural correlates of focused attention and cognitive monitoring in meditation. *Brain Res Bull, 82*(1-2), 46-56. doi:10.1016/j.brainresbull.2010.03.001

Miyoshi, T., Tanioka, K., Yamamoto, S., Yadohisa, H., Hiroyasu, T., & Hiwa, S. (2019). Revealing Changes in Brain Functional Networks Caused by Focused-Attention Meditation Using Tucker3 Clustering. *Front Hum Neurosci, 13*, 473. doi:10.3389/fnhum.2019.00473

Soni, R., & Muniyandi, M. (2019). Breath Rate Variability: A Novel Measure to Study the Meditation Effects. *Int J Yoga, 12*(1), 45-54. doi:10.4103/ijoy.IJOY_27_17

Van Overwalle, F., Baetens, K., Mariën, P., & Vandekerckhove, M. (2014). Social cognition and the cerebellum: A meta-analysis of over 350 fMRI studies. *Neuroimage, 86*, 554-572. doi:https://doi.org/10.1016/j.neuroimage.2013.09.033
